# Phytoplankton community structure and photophysiological strategies in the oligotrophic Indian Ocean spawning ground of Southern Bluefin Tuna

**DOI:** 10.1101/2025.09.18.677227

**Authors:** Joaquim Goes, Helga do R. Gomes, Xiwen Wu, Sven A. Kranz, Rasmus Swalethorp, Jinghui Wu, Anima Tirkey, Michael R. Landry

## Abstract

The Argo Basin, a persistently oligotrophic seascape downstream of the Indonesian Throughflow (ITF), is the only known spawning ground of Southern Bluefin Tuna (*Thunnus maccoyii*). Its surface waters are capped by the warm, low saline, nitrate-depleted Pacific Ocean outflow, that leads to intense water-column stratification and chronic nutrient scarcity. During the BLOOFINZ campaign (R/V *Roger Revelle*, January–March 2022), we combined high-resolution underway surveys with Lagrangian experiments to examine hydrography, nutrient stoichiometry, and phytoplankton photophysiology across the basin. This study was motivated by the need to: 1) identify the phytoplankton communities capable of persisting in this nutrient-impoverished region, and 2) uncover the unique photophysiological traits that underpin their survival and growth, particularly under acute nitrate limitation. Nutrient fields revealed strongly negative N*, positive Si*, and elevated Si:P ratios, indicating a system of acute nitrate scarcity but with residual phosphate and latent silicate availability. Phytoplankton communities were dominated by the cyanobacteria, *Prochlorococcus*, whose survival strategies included small functional absorption cross section of PSII (σPSII), enhanced PSII connectivity (*p*), highly elastic and reversible diel photophysiology that balanced high-light stress and nitrogen scarcity. Longer turnover times especially τ_2_ (µs; PSII–PSI electron transport) seen in the region around 115° and 120°E and at 15°S, and during Cycles 3 and 4 are indicative of populations that were being stressed by Fe limitation as well. Our inference of *Prochlorococcus* is supported by complimentary measurements of flow cytometry, phytoplankton pigment composition and microscopy all of which confirmed the dominance of *Prochlorococcus* in these nutrient impoverished waters. Our observations further indicate that mesoscale eddies generated by ITF–SICC (South Indian Counter Current) interactions, can episodically shoal the nutricline, triggering transient pulses of larger eukaryotes. Collectively, these results demonstrate how acute nitrate depletion, compounded by Fe stress, structures a microbial community finely tuned to oligotrophy and capable of sustaining a short food chain involving appendicularians that directly support larval tuna. The Argo Basin therefore exemplifies how circulation–nutrient coupling shapes microbial communities and ecosystem function within a globally significant spawning habitat.

**Highlights:**

1) Chronic nitrate limitation, with strongly negative N* and persistent N:P imbalance defines the Argo Basin while phosphate and silicate remain in relative excess.
2) Picocyanobacteria especially *Prochlorococcus* dominate the ecosystem, sustaining productivity under extreme oligotrophy through photophysiological strategies of reduced σPSII, enhanced PSII connectivity, and diel recovery from light stress.
3) Mesoscale eddies at the ITF–SICC front provide episodic relief, shoaling the nutricline and fueling transient pulses of larger phytoplankton that episodically reshape phytoplankton community structure and biomass
4) This oligotrophic–frontal seascape supports a short, efficient food web, enabling transfer of picophytoplankton to Southern Bluefin Tuna larvae via mainly appendicularians.

## 1. Introduction

Waters overlying the 5000-m deep Argo Abyssal Plain (hereafter, Argo Basin), a small sector of the southeastern Indian Ocean between northwestern Australia and Indonesia and immediately downstream of the Indonesian Throughflow (ITF), is a region of global oceanographic significance (Lee et al., 2015; Schneider, 1998; Sprintall and Révelard, 2014) This area represents the only low-latitude connection between major ocean basins, through which warm, low-salinity, relatively nutrient-rich water from the western Pacific is transported into the south tropical Indian Ocean. The hydrographic setting in this region is characterized by strong horizontal advection of warm, low-salinity, low-density Indonesian Through Flow (ITF) waters with a well-defined thermocline, weak vertical mixing, and a largely oligotrophic surface layer (Ayers et al., 2014; Condie and Andrewartha, 2008; Kehinde et al., 2023; Koch-Larrouy et al., 2015).

These conditions shape the phytoplankton structure (Furnas, 2007), physiology, and productivity (Kranz et al., this issue) that favor small, efficient primary producers. Phytoplankton communities in this region are primarily composed of pico-sized (<2 µm) organisms dominated by *Prochlorococcus* and some localized hotspots of picoeukaryotes (Selph et al., this issue-a; Yingling et al., this issue). Despite its ecological and oceanographic importance (Davis et al., 1990a; Davis et al., 1990b) the downstream ITF region remains sparsely studied. An earlier, extensive, study (Landry et al., 2022) on biomass structure, production and grazing rates from temperate to tropical waters, along the historic 110°E in the eastern Indian Ocean found *Prochlorococcus* dominated productivity, microzooplankton dominated grazing, with production and grazing tightly coupled and balanced. A notable gap is the lack of detailed studies on the survival strategies of phytoplankton, particularly their photosynthetic efficiencies, light-harvesting capacities, nutrient preferences, and growth dynamics. Understanding how stratification, light, and hydrodynamically driven nutrient fluxes, modulate the structure of phytoplankton communities and their photophysiology is essential for comparing this system among tropical–subtropical oceans (Behrenfeld et al., 2005; Wang et al., 2023). This is especially important because phytoplankton underpin the food web and influence higher trophic levels. In the Argo Basin, their role is amplified by supporting the survival and growth larvae of Southern Bluefin Tuna (SBT, *Thunnus maccoyii*), for which the basin is the only known spawning ground (Davis et al., 1990a; Davis et al., 1990b; Farley and Davis, 1998a).

From late January to early March 2022, the BLOOFINZ (Bluefin Larvae in Oligotrophic Ocean Food webs, Investigations of Nutrients to Zooplankton) campaign conducted an intensive end-to-end ecosystem survey and experimental study in the downstream ITF region (Landry et al., this issue-b). While the primary focus was food-web interactions affecting larval SBT feeding and growth, it provided a unique opportunity to examine phytoplankton dynamics in relation to hydrography, nutrient supply and grazing. Depth-resolved sampling (pigments, flow cytometry, microscopy; (Selph et al., this issue-b; Yingling et al., 2021) characterized community composition, while measurements of photosynthetic carbon fixation and nitrogen uptake assessed nutrient assimilation strategies and photosynthetic rates (Kranz et al., this issue; Yingling et al., this issue).

Here, we present underway measurements of high temporal-spatial resolution that provide a broad overview of phytoplankton photophysiology across the blue fin tuna spawning grounds, These measurements are complemented by 3–5-day Lagrangian experiments (Landry et al., this issue-b), which examine the photophysiology of resident phytoplankton in relation to diurnal changes in photosynthetically available radiation (PAR), as well as spatiotemporal variations in hydrography and nutrient availability. These measurements included the photophysiological signatures of light harvesting and electron transport (F_v_/F_m_, σPSII, connectivity *p*, and turnover times) as diagnostic indicators of growth strategies, nutrient and trace metal stress (Behrenfeld and Milligan, 2013; Gorbunov and Falkowski, 2020; Kolber and Falkowski, 1993; Oxborough et al., 2012; Schallenberg et al., 2020; Schuback et al., 2015; Schuback and Tortell, 2019; Schuback et al., 2021), and survival modes of pico-phytoplankton. By integrating these datasets, we sought to resolve the growth dynamics of small sized phytoplankton in a nutrient-limited tropical marine environments and to assess their ecological roles in the food web that supports SBT larvae.

## 2. Materials and Methods

### 2.1. Sampling design

The BLOOFINZ campaign, conducted aboard R/V *Roger Revelle* (cruise RR2201) from late January to early March 2022, involved an extensive suite of oceanographic observations from 12° to 17.1°S, 114 to 130°E, over the Argo Basin and adjacent Australian coastal margin (Landry et al. this issue, a) (Fig. 1). The study was planned to coincide with the peak SBT spawning season between January and February (Davis et al., 1990a; Davis et al., 1990b; Farley and Davis, 1998b).

**Figure 1.**
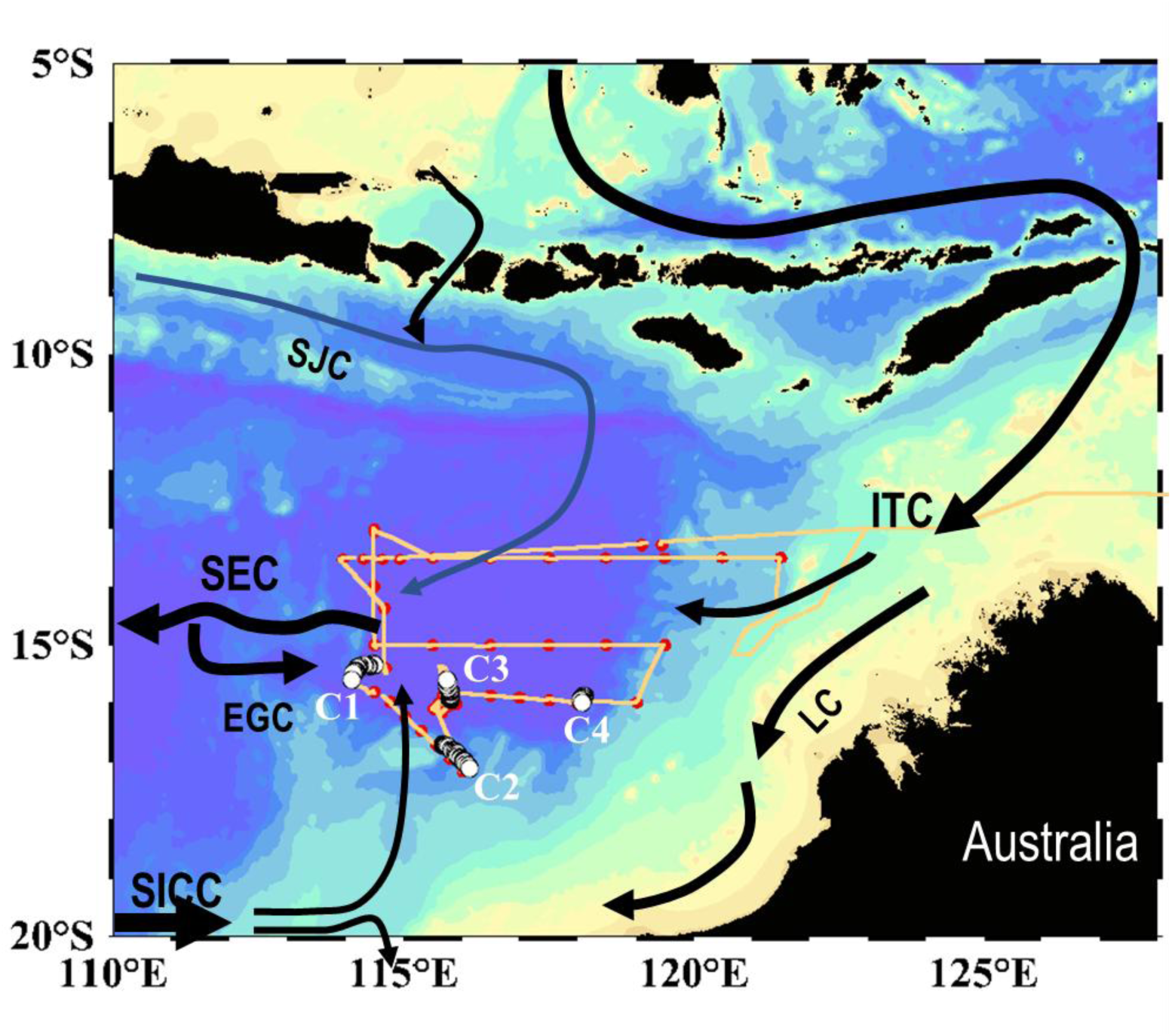
Map of southeastern Indian Ocean showing Indonesian Throughflow (ITF), South Equatorial (SEC), South Java (SJC), East Gyral (EGC) and Leeuwin (LC) currents. Orange line depicts continuous sampling along cruise track while red dots depict stations occupied. Cycle stations 1 to 4 are marked as C1 to C4 and denoted as white dots with black border.

Sampling on RR2201 was conducted in two modes: (1) underway sampling to capture mesoscale spatial variability of surface waters along the cruise track, and (2) point sampling, in which water from discrete euphotic-zone depths at predetermined locations was collected with a 12 × 10 L Sea-Bird^®^ CTD rosette. In the underway mode, several instruments were integrated into the ship’s uncontaminated seawater line that received water pumped with a diaphragm pump from ∼6 m depth at 4 L min ¹ into the ship’s wet lab and de-bubbled prior to delivery to each instrument.

The BLOOFINZ cruise began with underway and point sampling along an east-to-west transect along ∼13°S between 135° and 114.5 °E and across the Argo Basin (Fig. 1). This survey included the shallower regions of the Australian coastal margin, influenced by tidal activity and the open ocean. Upon completion of the east-west transect, the ship sailed south into the western Argo Basin, where the first of four Lagrangian process studies (hereafter “Cycles”) was initiated. The Cycle experiments followed a satellite-tracked drift array, allowing repeated sampling of the same water parcel and monitoring of phytoplankton community composition, biomass and photophysiology using several instruments described below. The Lagrangian studies followed a similar protocol to described in Landry et al. (2009) with measurements of stable isotopic composition, nutrient concentrations, nitrate uptake rate, N_2_ fixation, primary production, phytoplankton growth, micro- and mesozooplankton grazing, larval feeding and growth, and passive and active export (Stukel et al., this issue). Following the cycle experiments, two additional transects were surveyed across the basin: one at 15°S (east-to-west) and another at 13.5°S (west-to-east) to Darwin, Australia.

Nutrient samples from various depths, sampled using a Niskin rosette with CTD sensors were filtered through acid-cleaned 0.2-µm inline filter capsules (Pall-Gelman) into Nalgene HDPE bottles and immediately frozen at −20 °C. Nutrient analysis was done at the Scripps Institution of Oceanography Analytical Facility using a Seal AutoAnalyzer. Chlorophyll *a* (Chl *a*) concentrations at CTD sampling depths were determined onboard using a calibrated Turner Designs 10AU fluorometer (Holm-Hansen and Riemann., 1978). Samples were filtered onto GF/F filters and extracted overnight in 90% acetone at -20°C.

The in-line instruments data include Sea-Bird^®^ temperature and salinity sensors and a calibrated Turner Designs fluorometer for Chl *a* measurements. Photophysiological measurements were conducted with two fluorometers : (1) a Wet Labs^®^ Automated Laser Fluorescence Analyzer (ALFA; (Chekalyuk and Hafez, 2008; Chekalyuk et al., 2012; Goes et al., 2014a; Goes et al., 2014b), (2) a Rutgers University Fluorescence Induction and Relaxation (FIRe)^®^ instrument (Gorbunov and Falkowski, 2004; Gorbunov and Falkowski, 2020; Gorbunov et al., 2001). The instrument array also included a phytoplankton imaging system (Poulton, 2016), the Fluid Imaging Inc.^®^ FlowCAM which automatically detects, images, enumerates, and measures particles. Common measurements include the Equivalent spherical diameter (ESD), length, width, and aspect ratio, area and volume. Continuous measurements of phytoplankton fluorescence properties and images were obtained by pumping seawater (approx. 3 m depth sampling) from the ship’s flow-through system in darkness (all exposed tubes were covered with black tape) into the instruments’ sample chambers.

In addition to the continuous measurements, samples from various depths at the point stations were also collected in amber bottles and the samples were dark acclimated for 30 minutes to allow the photosystem to deoxidize, an important requirement to assess the photophysiological properties of the photoautotrophic community. Subsamples from these bottles were also used for analysis in the ALFA and in the FlowCAM, which have been described in more detail below.

### 2.2. Automated Laser Fluorescence Analyzer (ALFA)

The ALFA combines high-resolution spectral measurements of blue (405 nm) and green (532 nm) laser-stimulated fluorescence with spectral deconvolution to obtain fluorescence signals attributable to Chl *a*, phycoerythrin (PE), and colored dissolved organic matter (CDOM) (Chekalyuk and Hafez, 2008; Chekalyuk et al., 2012). All fluorescence values are Raman- normalized, with the exception of Chl *a*, which was calibrated with acetone extracted samples measured in a Turner Designs^®^ Trilogy Fluorometer (Holm-Hansen and Riemann., 1978).

Values for all other variables are expressed as relative fluorescence units (RFU). Blue-laser spectra allow estimates of Chl *a*, CDOM, and maximum PSII quantum yield (Fv/Fm; dimensionless), whereas green-laser spectra resolve three PE classes: PE-1 (565 nm; oligotrophic cyanobacteria with high PUB/PEB), PE-2 (578 nm; coastal-type cyanobacteria with low PUB/PEB), and PE-3 (590 nm; cryptophytes) (Chekalyuk and Hafez, 2008; Goes et al., 2014a; Goes et al., 2014b). PUB/PEB refers to the ratio of two light-absorbing chromophores, phycourobilin (PUB) and phycoerythrobilin (PEB), of marine cyanobacteria, and serves as an indicator of the cell’s adaptation to different light qualities in its environment. The instrument is set up to make these measurements approx. every minute.

### 2.3. Fluorescence Induction and Relaxation (FIRe)

The FIRe is a fast repetition rate fluorometer (Gorbunov and Falkowski, 2004) that measures dark-adapted PSII characteristics with a 450 nm diode to study photosynthetic light-harvesting, the photochemistry of PSII, and photosynthetic electron transport down to carbon fixation. FIRe can accurately measure fluorescence signals in samples as low as 0.01 mg m^−3^ of Ch *a* and is thus well suited for these oligotrophic waters.

Whether inline or in the laboratory, the FiRE protocol is as follows (Bibby et al., 2008; Silsbe and Gomes, 2022): Immediately after dark acclimation, the base fluorescence is measured under low actinic light (F_o_). A subsequent saturating blue (450nm central wavelength, 30nm bandwidth) light pulse is applied for a duration of 100 µs to close all reaction centers and achieve maximal fluorescence (F_m_). This change in fluorescence (F_v_ = F_m_-F_o_) is associated with absorption and utilization of light energy in photosynthesis and is normalized to F_m_ (F_v_/F_m_, dimensionless) to deduce the quantum efficiency of photochemistry in PSII. Functional absorption cross-sections of PSII (*σ*PSII, Å^2^ quanta^-1^) are derived from the rate at which fluorescence increases from F_o_ to F_m_. Following the termination of the saturating flash, the fluorescence yield relaxation is recorded, which reflects the kinetics of electron transport downstream of PSII. The kinetics of the rising and falling cellular fluorescence are calculated from the FIRe profiles based on the biophysical model of Kolber et al. (1998) to quantify the following photophysiological parameters: a) F_v_/F_m_ (dimensionless; maximum PSII quantum yield); b) σPSII (Å² quantum ¹; functional absorption cross-section); c) *p* (dimensionless; connectivity factor) ; d) turnover times τ_1_ (µs; QA re-oxidation); and e) τ_2_(µs; PSII–PSI electron transport). The FIRe was set up to obtain these measurements every 2 min.

F_v_/F_m_ for healthy, non-stressed open-ocean assemblages typically vary from ∼0.55 to 0.65. In *very* oligotrophic or nutrient-stressed layers these values are <0.5 (Falkowski and Kolber, 1995). A surface midday dip with recovery near dusk is typical photoprotection, not necessarily chronic stress ((Gorbunov and Falkowski, 2020). The FIRe, provides F_v_/F_m_ for a very short, saturating flash that excites the PSII only once and is called single turnover F_v_/F_m_ (ST). Conversely, when the light flashes are long enough to fully reduce the plastoquinone pool and close all the PSII reaction centers, the F_v_/F_m_ is referred to as multiple turnover F_v_/F_m_ (MT).

σPSII (Å^2^ quantum^-1^; functional absorption cross-section) reflects how much light a single PSII unit can capture. Larger σPSII values are a sign of photoacclimation to low light or nutrient stress, whereas smaller σPSII are indicative of more efficient, compact antenna systems, typical of high-light or nutrient-replete conditions.

*p* (dimensionless; connectivity factor), describes the probability that excitation energy absorbed by an antenna pigment can be shared between PSII reaction centers. In essence, *p* describes how absorbed photons are delivered to open reaction centers. So, if *p* approaches 0, it is indicative of no sharing of excitation energy; if *p* >0, it is indicative that the antenna systems are coupled and excitation energy can be redistributed efficiently among multiple PSII units.

Turnover time τ_1_(µs; QA re-oxidation) is the rate of primary electron transfer on the acceptor side of PSII. Short τ values indicate fast electron transfer, typical of healthy, nutrient-replete phytoplankton with well-functioning PSII reaction centers. Longer τ values indicate a slower re-oxidation of Q, which can signal stress or limitation especially nitrogen or iron both of which restricts synthesis of electron transport proteins and PSII repair machinery.

Turnover time τ_2_ (µs; PSII–PSI electron transport) reflects the rate of re-oxidation of the plastoquinone pool (Qb and beyond) and the reopening of PSII reaction centers after Q has transferred its electron. In other words, it captures the efficiency of electron transfer between PSII and PSI. Short τ values indicate fast plastoquinone turnover and efficient downstream electron flow rate associated with healthy, nutrient-replete cells where electron transport proteins and cofactors (e.g., iron-containing cytochromes) are abundant. Long τ values suggest a bottleneck further downstream of PSII, where electron carriers or cofactors are in short supply, symptomatic of nutrient, Fe stress or chronic light stress, where PSII damage is not fully repaired and the turnover slows.

These real-time, sensitive, non-destructive measurements provide instantaneous indicators of photosynthetic competency and can diagnose inorganic nutrient and iron limitation, photoacclimation, and photoinhibition (Behrenfeld and Milligan, 2013; Gorbunov and Falkowski, 2020; Schallenberg et al., 2020; Schuback et al., 2015).

### 2.4. Fluid Imaging FlowCAM

The FlowCAM imaging system (Poulton, 2016) has been used extensively to map phytoplankton communities ranging from 10–600 µm (Jenkins et al., 2016; Wu et al., 2022). In the present study, FlowCAM data obtained using a 4X flow cell, were used to distinguish broad phytoplankton functional groups, as well as the cell sizes and volumes that were converted to carbon contents using the carbon:volume ratios of Menden-Deuer and Lessard (2000). A notable limitation of the FlowCAM operated with a 4X flow cell, is that although it is able to count cells less than 10 µm, it cannot distinguish between *Prochlorococcus* spp. and *Synechococcus* spp.

## 3. Results

### 3.1 Hydrographic setting

Oceanographic conditions were oligotrophic, as evidenced by extremely low surface Chl *a* in the satellite composite of February 2022 (Fig. S1A). Sea surface temperature (SST) ranged from 26 to 30 °C (Fig. S1B), with warmer waters to the east under the influence of the ITF, which transports warm, low-salinity Pacific waters into the region, as observed in maps of sea surface height anomalies (SSHA) and surface currents (Fig. S1C and D). Waters to the west were colder, indicative of the South Indian Counter Current (SICC), an eastward-flowing current carrying cooler water into the warmer eastern boundary currents (Quadfasel et al., 1996). SSHA and surface current maps (Fig SIC-D), show that ITF–SICC interactions produce pronounced cyclonic and anticyclonic eddies, although cyclonic eddies did not appear intense enough to induce localized high Chl *a* patches visible from space (Fig. S1A).

A west-east section (Fig. 1a-d) along the 15.95 °S across the Argo Basin and prior to Cycle 1) shows uniformly warm (29–30 °C) surface waters, capping a mixed layer that varied from ∼5- 60 m (Fig. 2a-c). The salinity section (Fig. 2b) was marked by a low-salinity (∼34.4–34.6) lens characteristic of ITF waters, overlying a subsurface salinity maximum (∼34.7–34.9) centered at ∼20–60 m at several stations. Westward, the ITF lens thinned and the thermocline shoaled by several tens of meters. A subsurface salinity maximum (Fig. 2b) (∼34.8–34.9) at ∼115.75°E, marked the intrusion of SICC-ventilated subtropical thermocline water into the westward flowing ITF waters. The resultant east–west tilt of isotherms, isohalines, and isopycnals (doming to the west, deepening to the east) is the hallmark of a baroclinic front formed where buoyant ITF outflow meets the eastward flowing SICC near 12–14°S. Subtle undulations in the density field suggest mesoscale variability consistent with eddy activity generated by shear between the ITF and SICC, with intermittent doming due to cyclonic mesoscale eddies eroding the ITF cap (Brink et al., 2007).

**Figure 2.**
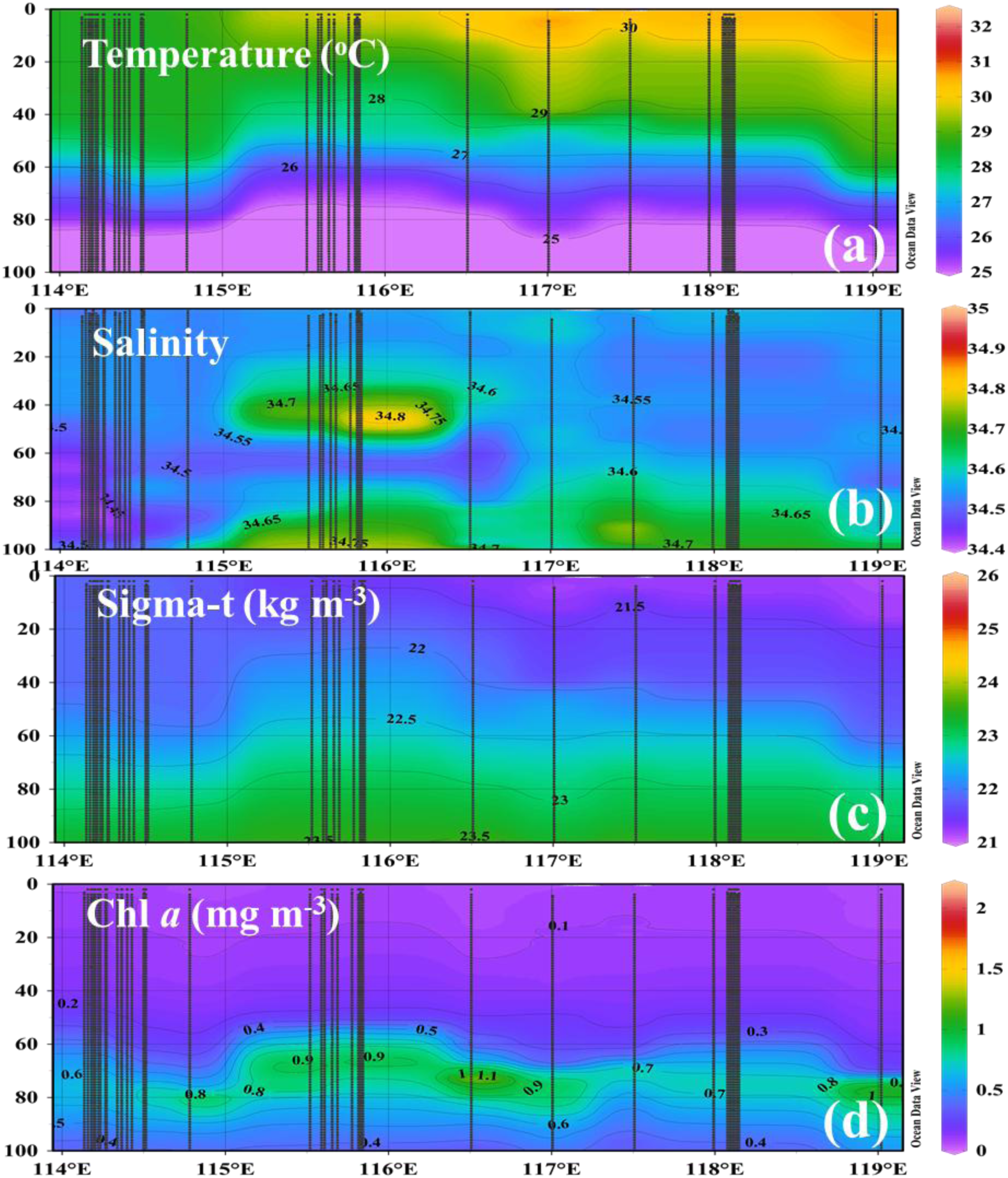
West-East section along 15.95°S across the Argo Basin showing a) Temperature (°C) b) Salinity (PSU), c) Density (kg m^-3^) and d) Chl *a* (mg m-3).

Surface Chl *a* concentrations were low (∼0.04–0.27 mg m^-3^), consistent with oligotrophic ITF waters (Figs. 2d and S1A). A deep chlorophyll maximum (DCM) was observed straddling the pycnocline, varying in depth from ∼60–90 m. In the region under the influence of the SICC where isopycnals, shoaled at ∼115.75°E, the DCM was shallower, more intense and Chl *a* concentrations high (∼0.7–1.1 mg m^-3^), reflecting uplift of the nutricline. The latter is defined as the depth at which nitrate concentrations first exceed 1 μM and is a proxy of nutrient supply to the upper mixed layer of the ocean. In the east, under the thick eastern ITF lens, the pycnocline deepened, and the DCM was correspondingly deeper and weaker.

### 3.2. Inorganic nutrients

Surface nutrients were undetectable at most stations in the upper 20 m; therefore, the plots in Fig. 2a–d begin at 20 m closer to the depth where tuna larvae were found. Nitrate (NO ; Fig. 3a) was nearly depleted basin-wide (generally <0.10 µM), indicating acute nitrogen limitation. The lowest values were measured over the southwestern/western area while slightly higher concentrations (∼0.08–0.12 µM) were seen in the north central and western regions where the thermocline shoaled. Episodic, modest injections of “new” nitrate that is rapidly consumed are associated with intermittent mesoscale doming and frontal mixing where the SICC impinges on the ITF front. Surface PO ^3^ concentrations (Fig. 3b) were also low (≈0.05–0.10 µM) but exhibited a more distinct west–east gradient than nitrate: higher to the west/south, lower toward the stratified east. This pattern reflects the hydrography, showing that cooler, saltier SICC- ventilated thermocline water brings nutrients to the western flank, while the warm, fresh ITF cap in the east suppresses vertical nutrient resupply. The smaller dynamic range of PO ³ relative to NO indicates a pronounced N:P imbalance, with NO the proximate limiting nutrient. SiOH ^4-^ was low in absolute terms (Fig. 3c) but increased toward the east, with a secondary enhancement toward the southwest. Low values prevailed over the northwest/central-west. The eastward rise is consistent with the ITF biogeochemical imprint (elevated Si:N and Si:P), reflecting upstream dissolution/denitrification and perhaps weak drawdown by diatoms or radiolarians. The southwest tongue likely reflects mesoscale upwelling or lateral intrusions from SICC, whereas mixing with subtropical surface waters on the western side dilutes SiOH ^4-^.

**Figure 3.**
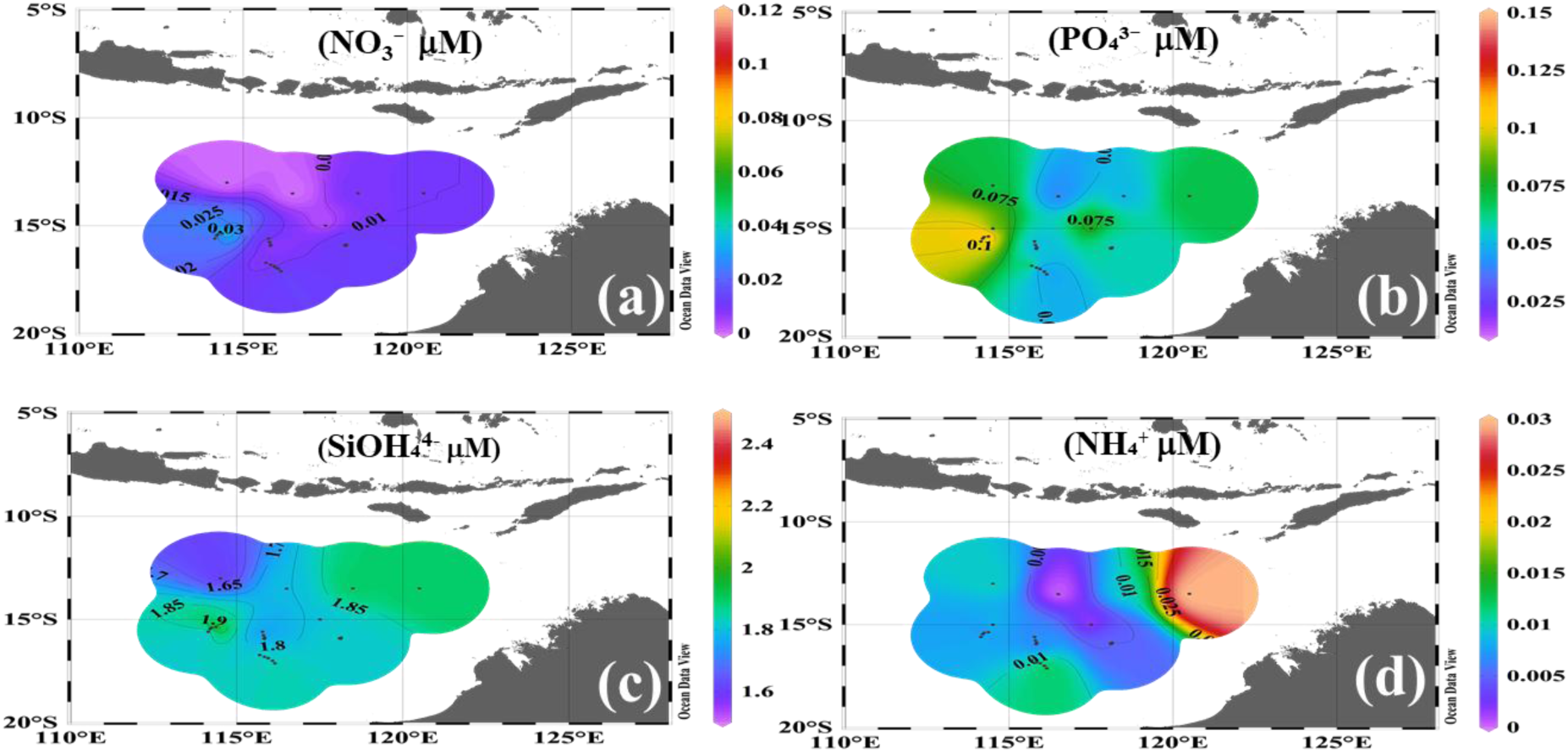
Spatial distribution of a) Nitrate (NO3^-^,μM), b) Phosphate (PO_4_^4-^ μM), c) Silicate (SiO_4_^4-^, μM) and d) Ammonium (NH^+^ μM) at 20m in the Argo Basin. Measurements were made in samples from the CTD rosette at point stations

NH showed a pronounced eastward maximum (∼0.10–0.15 µM) (Fig. 3d), where remineralization and zooplankton excretion supply regenerated N while euphotic-zone nitrification is suppressed. NH concentrations were lower westward, possibly due to stronger mixing and intrusions of the SICC that favor oxidation of NH to NO /NO .

At 60 m (Fig. S2a-d), nutrient distributions were different from those at 20m. The highest nutrient concentrations were along the east, where both NO and PO ³ attained maximum concentrations, with elevated concentrations extending to the central region and minima towards the west (Fig. 2a-b). This reversal relative to the surface reflects tidal mixing of the ITF waters in the shelf region. SiOH ^4-^ was likewise enhanced toward the east, with a secondary tongue toward the southwest and lows in the northwest/central-west, consistent influence of the ITF in the east and the SICC in the west (Fig. 2c). NH were high in the shelf region to the east indicating intense in-situ regeneration deeper within the euphotic zone and retained by frontal convergence, NH were also elevated in the northwestern part of the survey line (Fig. S2d).

N* values which indicates surplus or deficit of nitrate relative to phosphate were negative across the study area (Fig. 4a), with the most negative values in the warm, fresh ITF lens and along its downstream pathway corroborating the basin-wide nitrogen deficit relative to phosphate described above. N* was less negative at frontal/western margins when deeper water intrusions intermittently injected NO into the euphotic zone. However, N* did not approach zero, so nitrate limitation remained dominant. Si* was positive towards the east and near the front (Fig. 4b). High Si:N towards the ITF likely reflects rapid nitrate consumption relative to Si.

**Figure 4.**
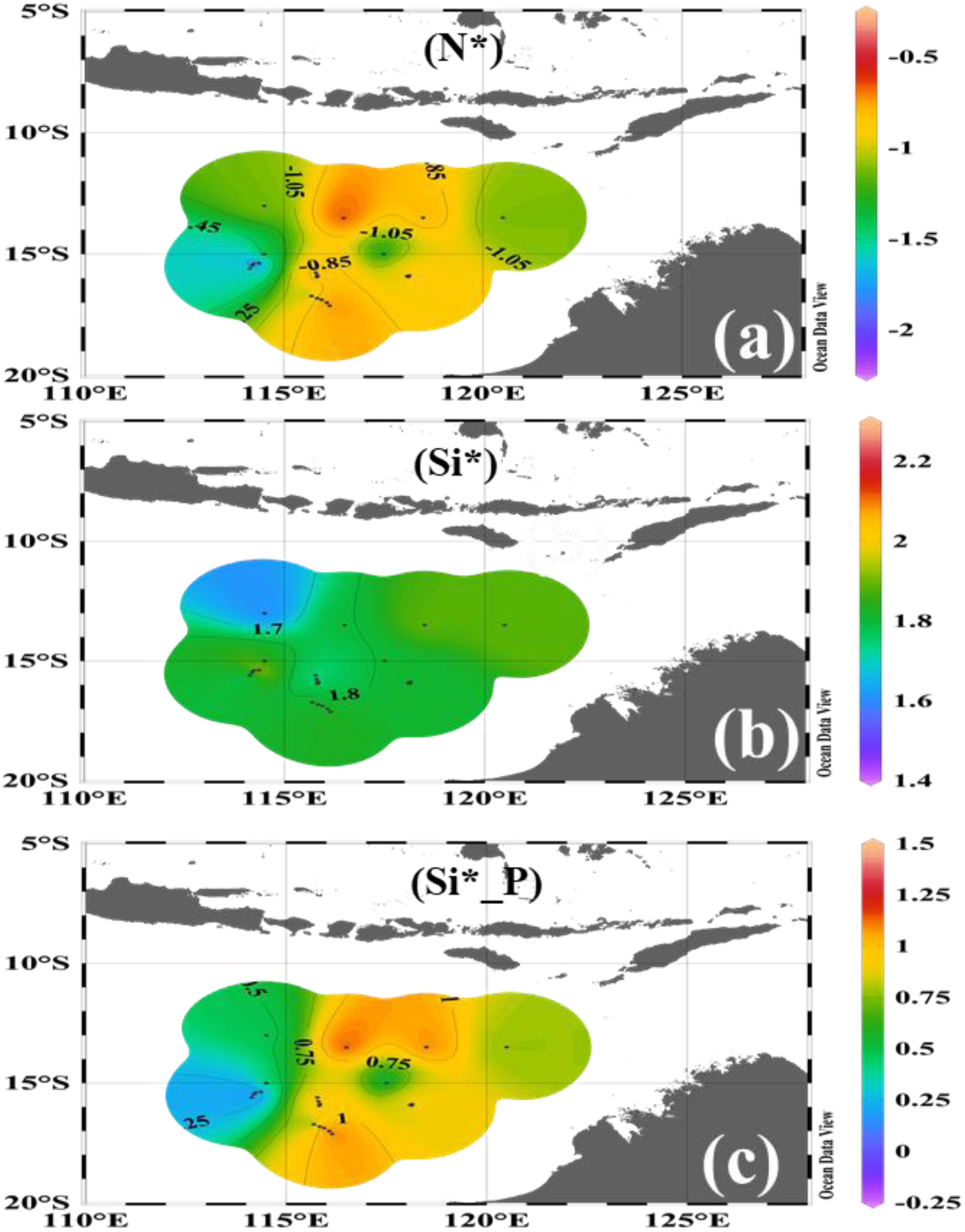
Spatial distribution of a) N*, b) Si* and c) Si:P at 20m in the Argo Basin. Other details as in Fig. 3.

At 60 m depth, the nutrient stoichiometry fields revealed distinct patterns, in all probability shaped by the mixing of ITF waters with the surrounding Indian Ocean. The N* distribution (Fig. S 3a) shows predominantly negative values, indicating nitrogen depletion relative to phosphorus even at depth, consistent with the inflow of ITF waters that are typically N-deficient. Localized patches of near-zero or slightly positive N* suggest areas where mixing with ambient Indian Ocean waters partially offsets the nitrogen deficit. The Si* field (Fig, S3b) highlights a relative excess of silicate over nitrate, with positive values extending across much of the section. The highest Si* values occur toward the central and eastern portions of the basin, consistent with silicate-rich ITF inputs. In contrast, the Si:P stoichiometric ratio (Fig. S3c) shows elevated values across much of the study area, exceeding Redfield proportions and further emphasizing the strong silicate signal delivered by the ITF. The highest Si:P values are concentrated toward the south and southeast, aligning with areas where ITF inflow mixes with the SICC. Consistent with the distribution of nutrients at 20m, this pattern suggests that the vertical and lateral transport of ITF waters not only imprints the nutrient fields but also sets up stoichiometric imbalances that may shape phytoplankton community structure.

### 3.3. High-resolution hydrography, phytoplankton groups and photophysiological measurements

High-resolution shipboard SST and SSS measurements (Fig. 5a-b) are consistent with satellite observations (Fig S1a-b) as well as the point sampling of Figs. 2a-d, revealing a strong east–west gradient along the cruise track, with the fresh water ITF lens to the east and the saltier SICC-influenced waters to the west.

**Figure 5.**
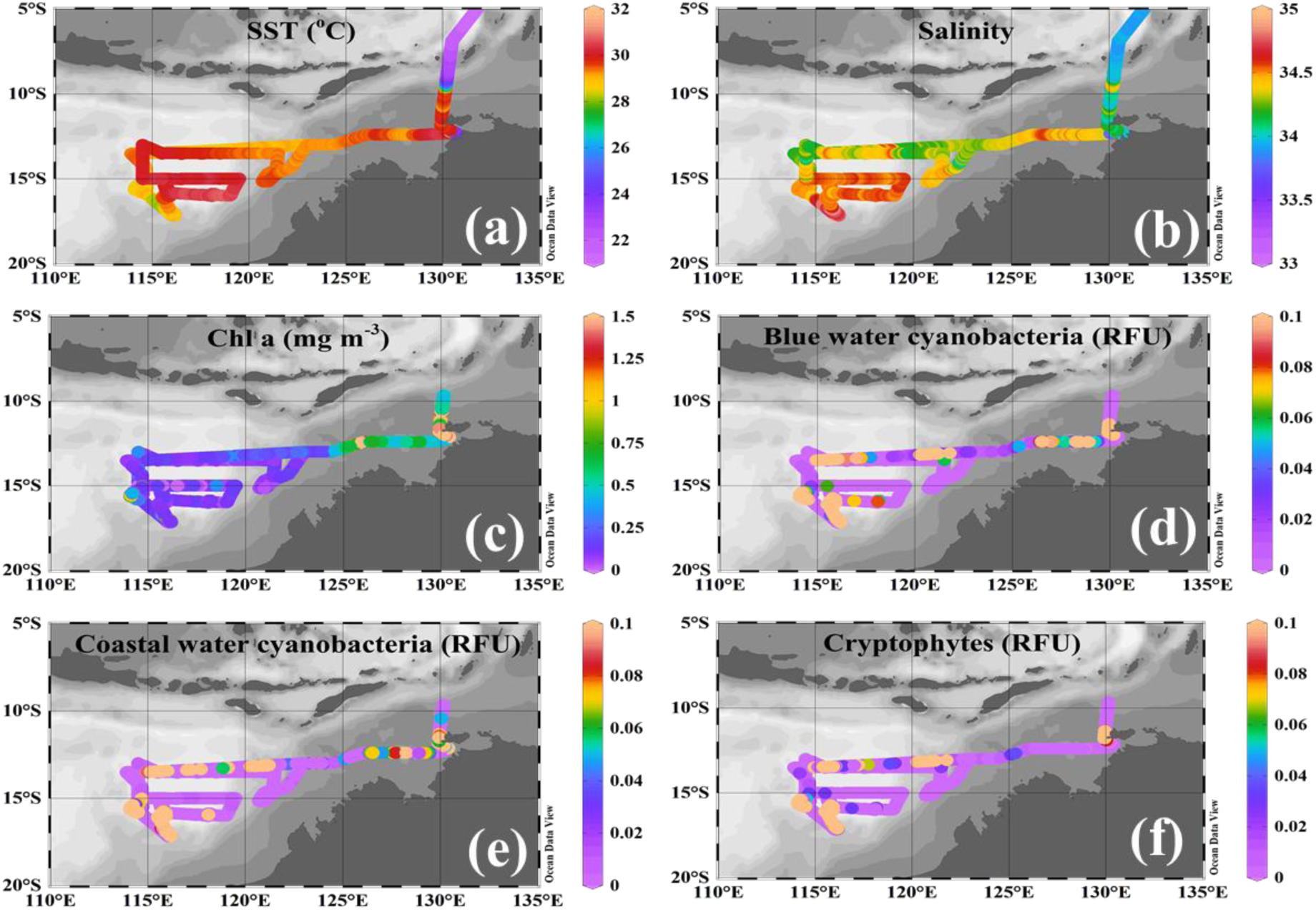
Distribution of a) SST(°C), b) Salinity (PSU), c) Chl *a* (mg m^-3^), d) Blue water cyanobacteria (RFU), e) Coastal water cyanobacteria (RFU) and f) Cryptophytes (RFU) measured by the ALFA attached to the continuous flowthrough system.

Surface Chl *a* obtained from the high resolution ALFA measurements was uniformly low (Fig. 5c), along the northernmost transect but modest increases were observed east of ∼125°E, a region that experiences tidal mixing (Brink et al. 2007; Schiller, 2011). Unlike in the Amazon river plume (Goes et al., 2014b) and the Bering Sea (Goes et al., 2014a), on account of the low biomass, ALFA fluorescence signatures were not strong enough to clearly differentiate between blue water and coastal cyanobacteria. Co-located hot spots of both types of both coastal water and open ocean cyanobacteria were observed along the 14°S transect (Fig. 5d-e), as well as elsewhere in the study area. There was some localized enhancement of coastal cyanobacteria near the shelf where the overlying waters are influenced by coastal processes including tidal mixing (Fig. 5e). The general distribution is consistent with a picocyanobacterial assemblage adapted to strong stratification and severe nitrate depletion. Cryptophytes (Fig. 5f) were generally sparse in the oligotrophic core but also formed distinct hotspots, co-located with the strongest hydrographic gradients and where 60-m NO /PO ³ /Si(OH) maxima impinged on the euphotic zone. Their distribution indicates sensitivity to episodic nutrient–mixing events as these features were not seen along the eastward transect that was undertaken after approximately a month later after completing the cycle experiments, indicating the episodic nature of these hotspots.

Both estimates of photochemical efficiency (F /F and F /F (MT) reached maxima on the eastern frontal limb and closer to the shelf region (≈0.53–0.58) indicating healthy populations. Away from the shelf region, values decreased to ≈0.44–0.50 over the western/southwestern loop and were lowest between 115° and 119°E, 15° and 13°S (Fig. 6a, d).

**Figure 6.**
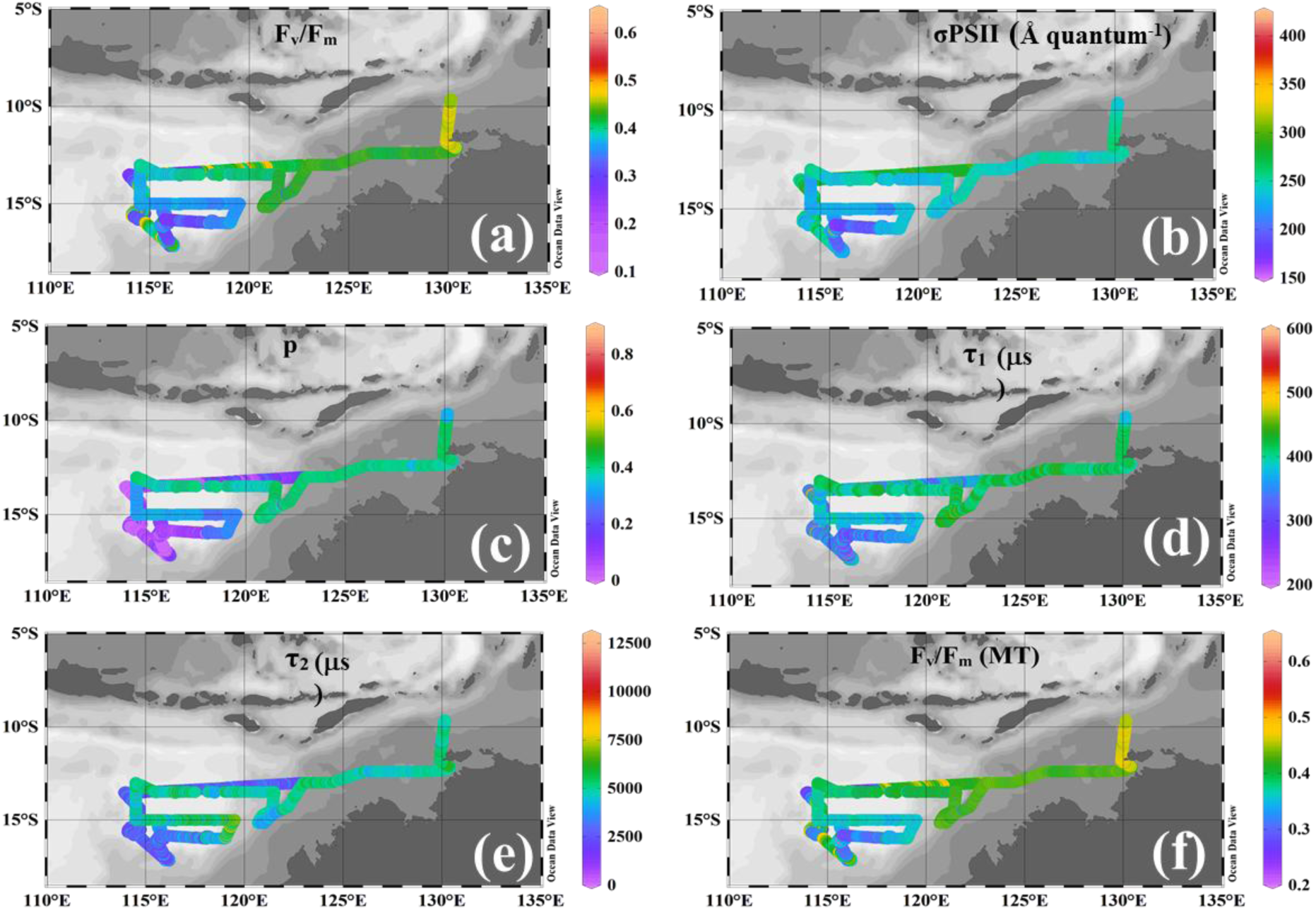
Distribution of variable fluorescence properties a) F_v_/F_m_ (ST, (maximum PSII quantum yield; dimensionless) b) σPSII (functional absorption cross-section; Å^2^ quantum^-1^;), c) *p* (connectivity factor; dimensionless;) d) turnover times τ_1_ (QA⁻ re-oxidation; µs) e) τ_2_ (PSII–PSI electron transport; µs) and f) F_v_/F_m_ (MT) by the fast repetition rate variable fluorometer FIRe attached to a continuous flowthrough system.

Both the effective PSII absorption cross-section (σPSII) (Fig. 6b) and the PSII connectivity parameter (*p*) (Fig. 6c) followed a similar pattern. σPSII transitioned from ∼170–230 in the west to 250–320 Å quantum ¹ eastward (Fig. 6c), and PSII connectivity parameter, *p* transitioned from low in the west to moderate–high in the east (Fig 6d). Time constants of reoxidation (τ, fast closure/primary charge-separation; τ, slower QA re-oxidation/RC reopening) were shorter in the west and southwest and longer in the east (τ ≈350–550 → 600–800 μs; τ ≈400– 700 → 800–1200 μs) (Fig. 6e-f). Thus, the eastern sector is marked by higher F /F, larger σPSII, greater PSII connectivity, and a slower turnover, while the west is characterized by lower F /F, smaller σPSII, weaker connectivity, and faster turnover. The photophysiological signatures of low F /F and σPSII and shorter τ’s are consistent with physiological stress under acute nitrate limitation (smaller effective antennae, reduced connectivity, faster RCII turnover). There is also the possibility that SICC/eddy mixing exposes cells to higher instantaneous light.

### 3.4. Cycle photophysiological observations

During Cycle 1 from 4^th^-8^th^ Feb. 2022 (Figs. 7a-g) which followed a storm, PAR showed diurnal cycles with mid-day peaks >2,000 µmol m ² s ¹. Clouds prevailed over the first three days with the sun appearing intermittently as is seen in the variability in PAR over these days. Concomitant with this, F_v_/F_m_, σPSII and connectivity factor (*p)* also showed distinct diurnal variations. Because F_v_/F_m_ was measured following dark acclimation in the FIRe, the observed changes most likely reflect photoinhibitory damage to PSII reaction centers (qI), rather than non-photochemical quenching processes associated with energy dissipation (qE) or state transitions (qT). Overnight values of F_v_/F_m_ (0.46–0.50) indicate partial recovery of PSII efficiency in the absence of light stress, with a steady increase into early morning as cells re-established photochemical capacity. The subsequent decline to midday minima (∼0.33–0.36) points to the accumulation of photodamage under peak irradiance. This interpretation is consistent with the concurrent increase in connectivity (*p*) during early morning (∼0.12–0.20), which may reflect enhanced functional coupling among PSII units to optimize electron transport as light levels rise. The relaxation of *p* at night suggests that this plastic response is reversible and linked to daily light–dark cycling, further highlighting the role of pico-phytoplankton in balancing photoprotection with efficient light harvesting under nutrient-limited tropical conditions. Acute cloud cover on the first 3 days increased the maxima in F_v_/F_m_ as well as *p* suggesting that phytoplankton cells in the latter days (when cloud cover disappeared), were photoinhibited. σPSII showed smaller diurnal variations (average ∼250 Å quantum ¹) with modest midday declines.

**Figure 7.**
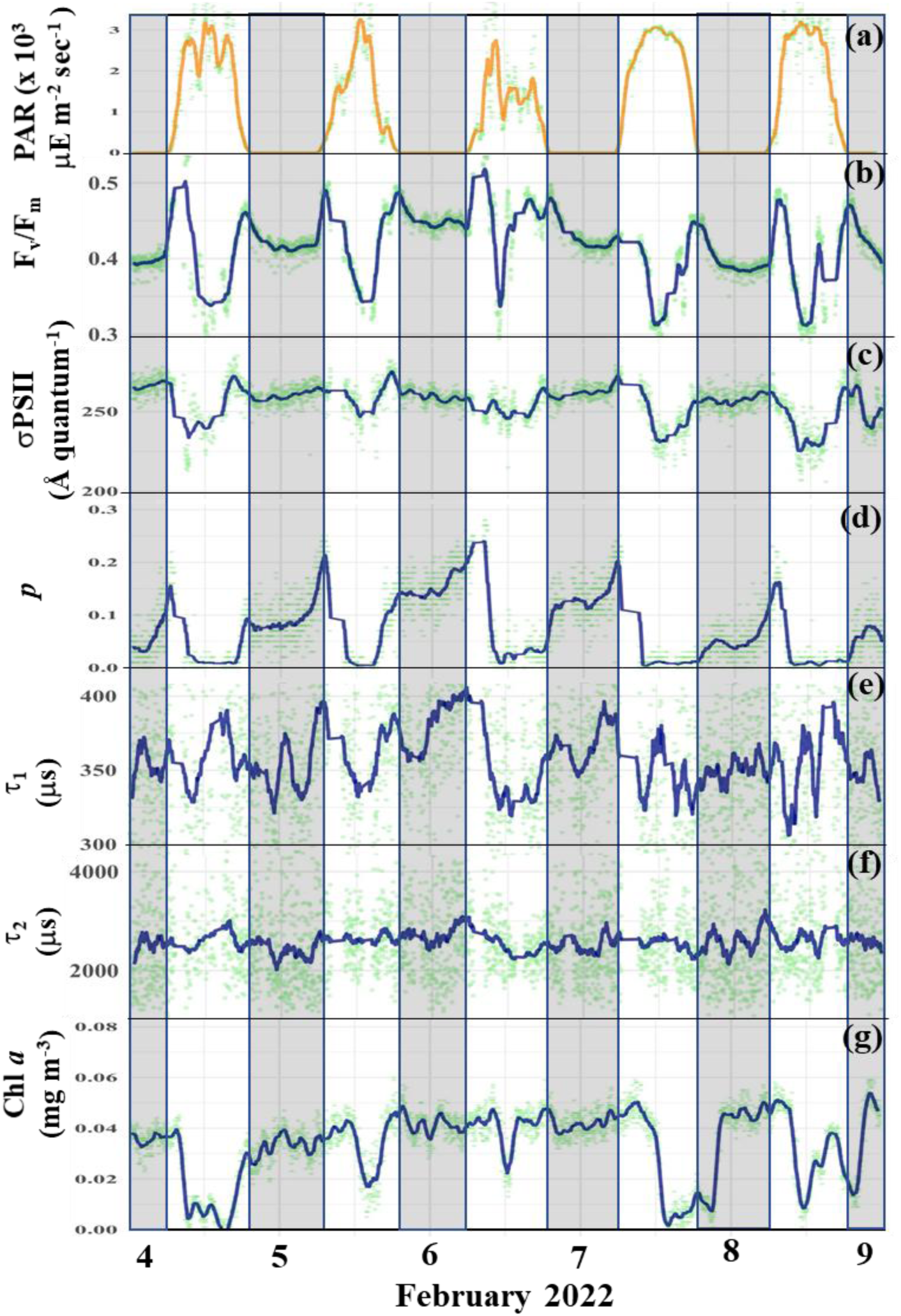
Diurnal variability in a) PAR, b) F_v_/F_m_, c) σPSII, d) *p,* e) τ_2_, f) τ_2_ and g) Chl *a* during Cycle 1. Dark columns denote night with the middle of column indicating midnight.

During the day both τ and τ increased, indicating slower electron turnover. τ rose from its nocturnal baseline (∼300–330) to 350–400, while τ showed a larger amplitude increase, from ∼2000–2500 at night to ∼3000–3800 during the day. This daytime slowing of electron turnover likely reflects the buildup of excitation pressure as photosynthetic electron transport is driven by incoming photons. The increased turnover times are consistent with heightened demand on downstream acceptors and the activation of regulatory processes such as non-photochemical quenching and cyclic electron flow, which act to balance energy input with metabolic capacity. During the night, both τ and τ decreased and stabilized, with τ around 300–330 and τ between 2000–2500. The suppression of turnover times in darkness suggests more rapid electron relaxation when photochemical demand is absent, consistent with an open and relaxed photosynthetic apparatus. The lower variability at night further indicates that electron pools are in a more stable state, unperturbed by fluctuations in irradiance or photosynthetic regulation.

In addition to these day–night contrasts, both τ and τ exhibited subtle but reproducible increases midway through the night and again in the hours preceding dawn. For τ, these increases were modest (∼10–20 units above baseline), whereas τ rose by several hundred units. Although smaller in magnitude than the daytime peaks, these nocturnal and pre-dawn elevations suggest that electron turnover dynamics are shaped not only by instantaneous irradiance but also by slower, endogenous processes. Such patterns may reflect circadian regulation, metabolic rebalancing of stromal redox states, or pre-illumination adjustments that prepare the photosynthetic apparatus for the transition into the light period.

Chl *a* fluorescence was highest in darkness (∼0.03–0.04) and strongly quenched by day, steadily recovering thereafter until nightfall when it declined again.

Taken together, these results demonstrate that electron turnover times are tightly coupled to diel light cycles, with daytime illumination slowing both fast and slow electron pools, and darkness restoring rapid, stable turnover, yet also reveal subtle nighttime regulation that points to circadian or metabolic control of photosynthetic electron transport beyond the direct influence of light.

During Cycle 2 (10–15 February), the photosynthetic variables continued to exhibit strong light–dark dependence, though with some differences in magnitude compared to Cycle 1. Both F_v_/F_m_, and σPSII were consistently lower overall, suggesting reduced maximum photochemical efficiency and smaller functional antenna size during this cycle. Despite these reductions in absolute values, the diel pattern was more clearly expressed across all parameters. During daytime (light periods, white sections), F_v_/F_m_ declined further, while σPSII showed daytime depressions consistent with enhanced energy dissipation under high light. The parameter *p* also exhibited a pronounced increase under illumination, mirroring the light forcing in panel (a).

Similarly, both electron turnover times were elevated: τ rose from ∼300–330 at night to ∼360– 400, while τ increased more strongly from ∼2000–2300 in darkness to ∼3200–3700 in daylight, again accompanied by higher variability during the light period. During the night, all parameters converged towards stable baselines. F_v_/F_m_, recovered, σPSII increased, and both τ and τ declined to lower, more consistent values. In Cycle 2, this contrast was sharper than in Cycle 1: τ consistently suppressed at night to ∼300–320, while τ stabilized at ∼2000–2400, with minimal scatter.

Together, these results highlight that while Cycle 2 was characterized by somewhat lower F_v_/F_m_, and σPSII values overall, the diel structure was more clearly defined across the full set of photosynthetic parameters as compared to Cycle 1. This stronger alignment with the light–dark cycles suggests that electron turnover (τ, τ) and energy fluxes (p, σPSII) were more tightly synchronized with irradiance forcing during this cycle, despite the reduced baseline efficiency.

During Cycle 3 (16 to 20 February), the continuation of water masses from Cycle 1, (Figs. 9a-g), PAR was consistently high (2,500–3,000+ µmol m ² s ¹) under clear skies. all photosynthetic parameters continued to track the light–dark transitions, but their absolute values were much lower compared to Cycles 1 and 2. Predawn/evening F /F ∼0.35–0.37 dipped slightly to ∼0.32–0.34 at noon and recovered each evening, indicating reversible down- regulation. Similarly, σPSII was lower than at the previous Cycles, suggesting a diminished functional antenna size and photochemical efficiency under the prevailing conditions, but exhibited a diel rhythm. The parameter *p* also showed lower absolute values but, importantly, exhibited a much clearer diel rhythm, tightly aligned with the light forcing in panel. Both electron turnover times, τ and τ displayed well-ordered day–night cycles. During daytime, τ rose toward **∼**350–380, and τ toward ∼3000–3600, with enhanced variability under illumination. At night, τ and τ consistently decreased to **∼**300–320 and ∼2000–2400, respectively, with reduced scatter. Compared to Cycles 1 and 2, the diel modulation of both τ and τ was sharper, indicating stronger coupling to irradiance. Chl *a* concentrations were much lower during this Cycle and diel cycles were also much shallower compared to Cycles 1 or 2. This muted response suggests that the photosynthetic apparatus was operating under sustained stress, where baseline efficiency was already suppressed.

**Figure 8.**
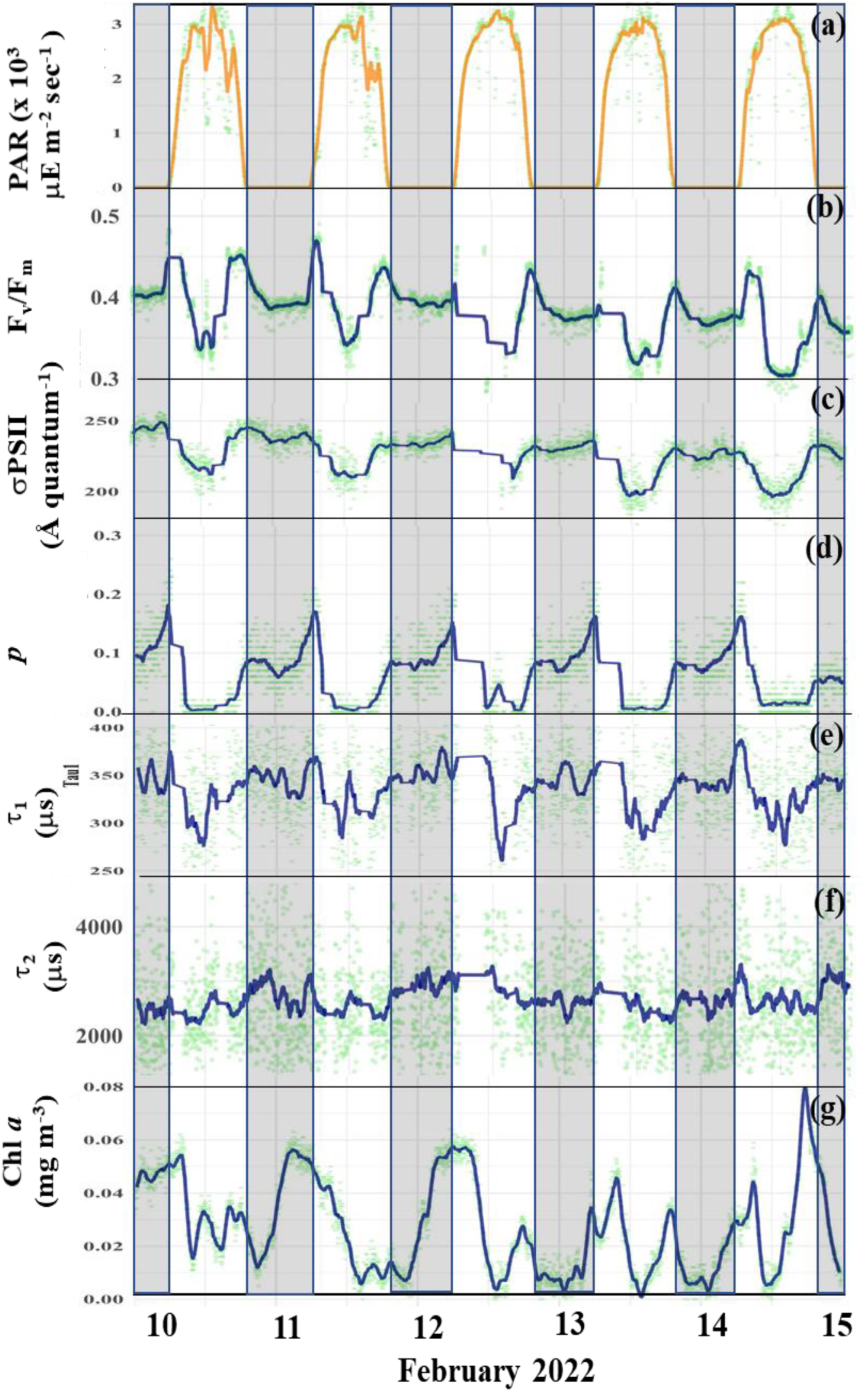
Same as Fig 7a-g but during Cycle 2

**Figure 9.**
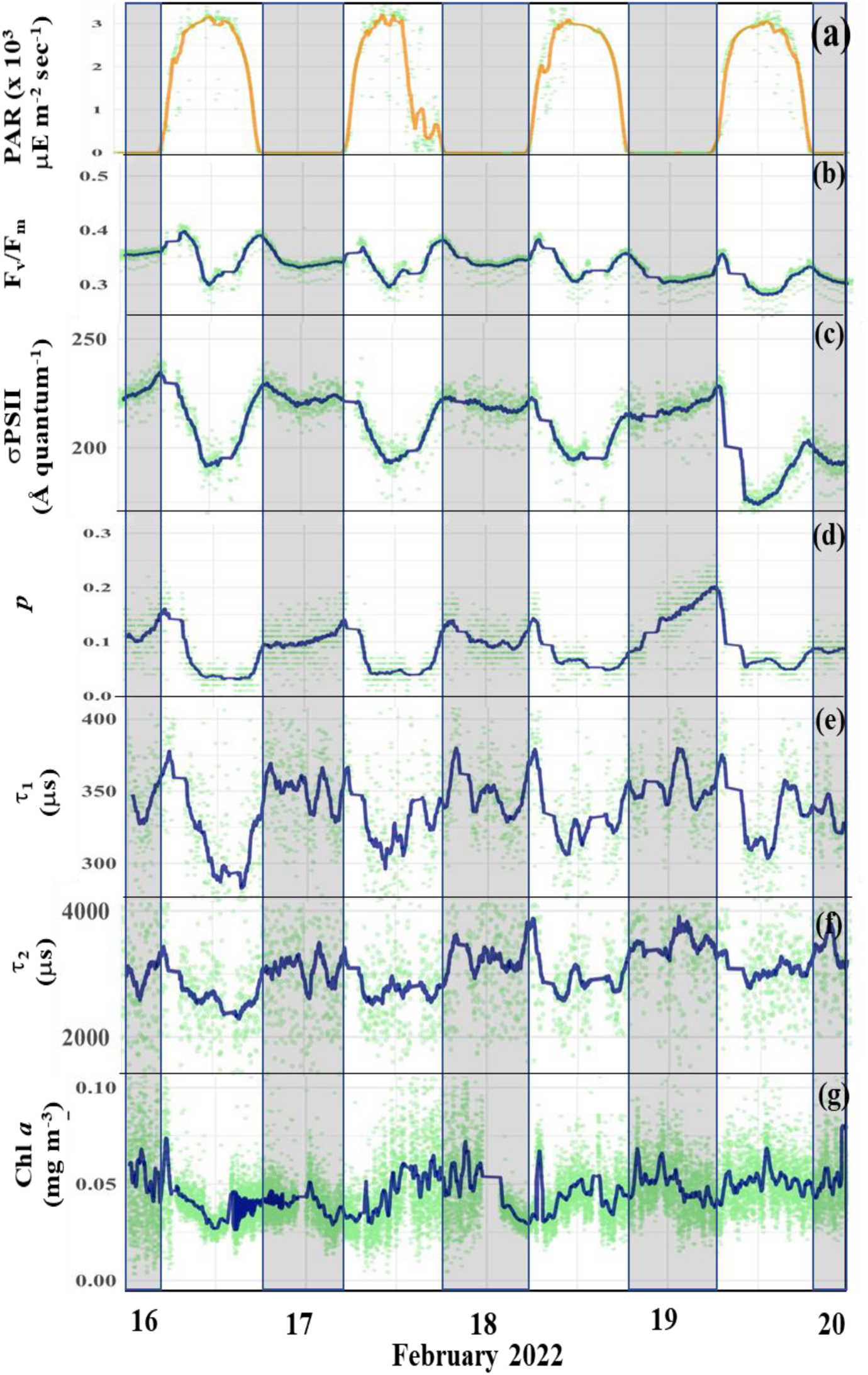
Same as Fig 7a-g but during Cycle 3

Cycle 4 (Fig. 10a-g), from 23–25 February showed slightly different patterns from the other 3 cycles. Mid-day PAR reached ∼2600–3100 µmol m ² s ¹. Diurnal changes in photophysiological properties were observed across all parameters but were much muted compared to the other cycles. Predawn/evening F /F ∼0.35–0.37 dipped slightly to ∼0.32–0.34 at noon and recovered each evening, indicating reversible down-regulation. Interestingly a major difference from the prior cycles was a slow but distinct increase in σPSII (morning ∼210–230; noon ∼195–210; evening ∼215–240 Å quantum ¹) from the 23^rd^, more so in *p* which rose sharply (∼0.06–0.08 to ∼0.22–0.26) and remained high. Both electron turnover times, τ and τ, displayed day–night alternation, with elevated daytime values (τ : ∼340–380; τ : ∼3000–5000) and suppressed nighttime baselines (τ : ∼300–320; τ : ∼2000–3000). What was different, from the previous Cycles, the daytime maxima of both τ and τ increased steadily across the 4 to 5- day sequence, producing a stepwise rise in turnover times rather than the repeating diel plateaus seen in earlier cycles. This increase albeit not so distinct was also observed in surface Chl *a* (∼0.02–0.03 → 0.04–0.05 mg m ³) with shallow midday troughs and little nocturnal decline.

**Figure 10.**
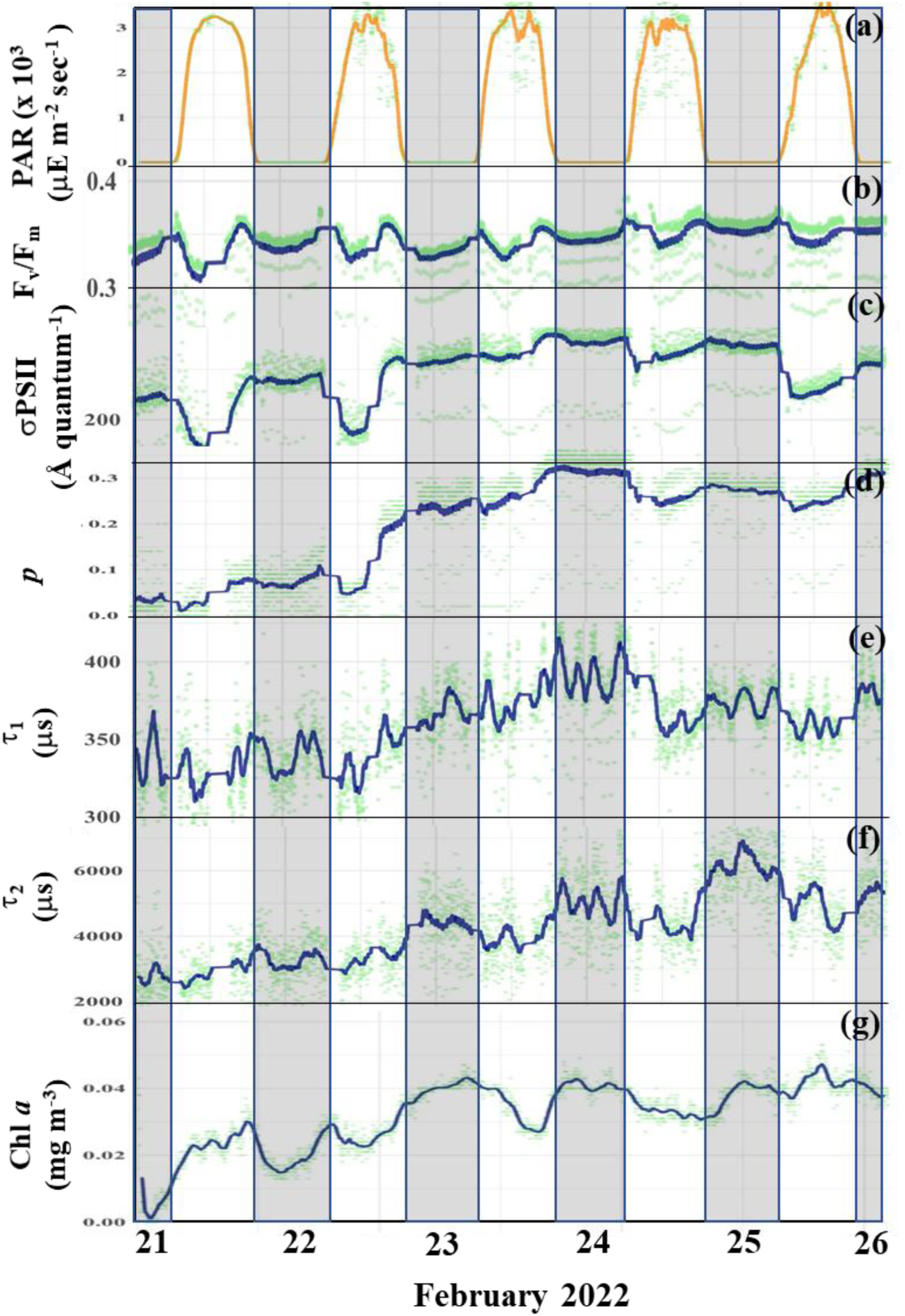
Same as Fig 7a-g but during Cycle 4

The progressive upward drift in baseline values in all photophysiological parameters, across consecutive days, represents a key distinction from the relatively stationary behavior observed in Cycles 1–3. This differing photophysiology in Cycle 4 was also noted by Kranz et al. (this issue) who measured the lowest variable fluorescence based Gross Primary Productivity (GPP) at this site. Based on a decreased Fv/Fm (0.23) but enhanced antenna size (σ) as well as the parameters of the Photosynthesis-Irradiance curves, they concluded that the photophysiological response of cells was in transitional state of high-light stress and acclimation, suggestive of mild photoinhibition, and incomplete acclimation to higher light. The larger σ suggests a partial enhancement of light-harvesting complexes, possibly reflecting improved nutrient availability or community shifts. Overall, they suggested cells were maintain efficient photosynthetic operation under fluctuating light and nutrient regimes.

### 3.5. Phytoplankton community structure

FlowCAM data revealed overwhelming dominance of picoplankton across all four cycles, reflecting the oligotrophic, stratified conditions typical of this subtropical gyre. These small cells represented the bulk of numerical abundance (often >80–90% of counts) (Figs. 11a-d). However, the intermittent appearance of larger eukaryotes - particularly dinoflagellates and diatoms, contributed substantially to carbon biomass in spite of their low abundance especially in Cycles 1 & 4 (Figs. 12a-d).

**Figure 11.**
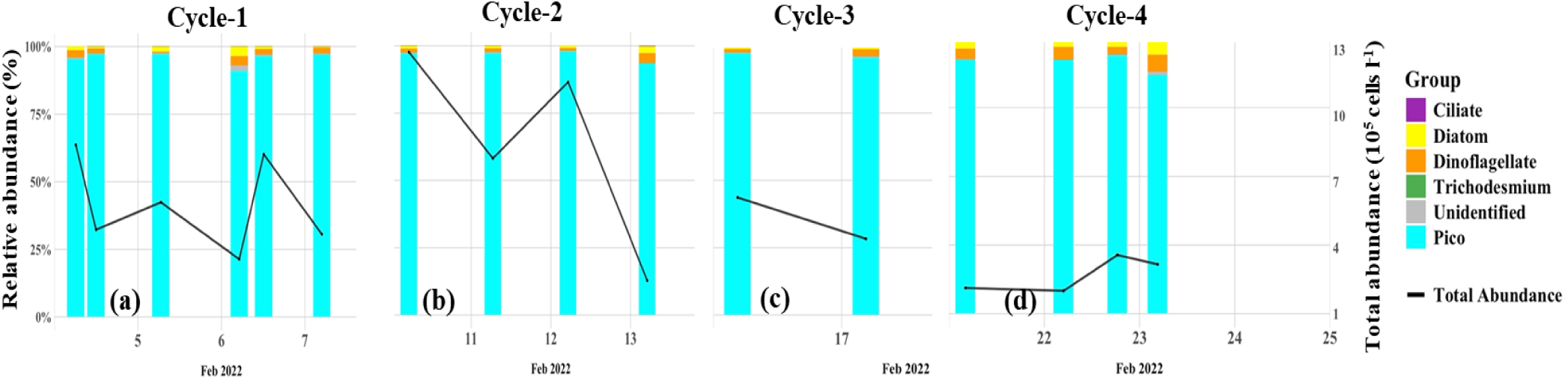
FlowCAM-derived percent phytoplankton community structure (primary axis) and total cell counts (cells L^-1^, secondary)

**Figure 12.**
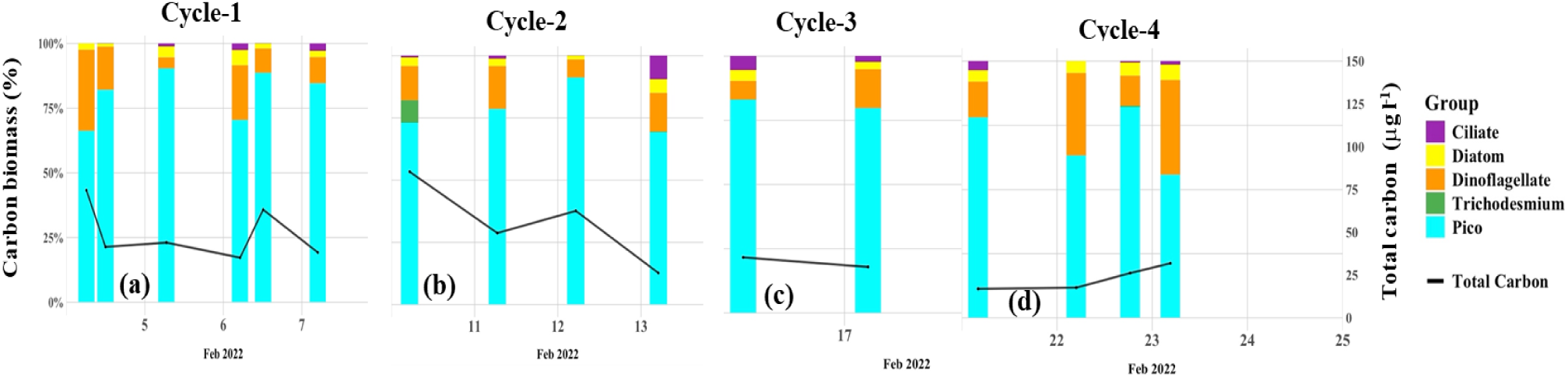
Changes in phytoplankton biomass (μg C l^-1^) for each group during the 4 cycles.

## 4. Discussion

### 4.1. Argo Basin hydrographic environment

The Argo Basin, the downstream offshore edge of the ITF in the southern Indian Ocean, is a hydrographically distinct region where warm, relatively fresh, nutrient-poor Pacific waters enter the Indian Ocean through narrow Indonesian island passages. As these waters spread westward, they help establish strong stratification (Atmadipoera et al., 2022; Ayers et al., 2014; Capotondi et al., 2012; Kehinde et al., 2023; Koch-Larrouy et al., 2015). Our study area was characterized by a shallow mixed layer, reinforced by surface heating and weak vertical exchange (Figs. 2a-c). These conditions remain for most of the year (Ayers et al., 2014), creating a persistently oligotrophic surface environment in which macronutrients (nitrate, phosphate, and sometimes silicate) are near or below detection limits and external supply occurs during rare episodic perturbations. To the west, mesoscale eddies generated by local baroclinic instabilities are the primary mechanism for intermittent nutrient injection into the euphotic zone (Figs. 2a-c) (Kehinde et al., 2023).

Throughout the upper mixed layer, N* was strongly negative, confirming the pervasive NO deficit relative to PO ³ . Chronic NO depletion is driven by the upstream influence of already nitrogen-depleted ITF waters and high biological demand in the stratified surface ocean. Experimental quantification of nutrient inventories, primary productivity and N_2_ fixation by Kranz et al. (this issue) revealed that N_2_ fixation provided a consistent new nitrogen source, contributing ∼16% to net primary production in the upper euphotic zone and indicating that N_2_ fixation supports most of the export production with recycling processes sustaining the rest of the community’s N demand. Positive Si* indicated silicate in excess of nitrate throughout the basin. Although SiOH ^4-^. is modest in absolute terms, it was sufficient relative to the near-absent NO pool indicating that absence of diatoms is primarily from nitrogen deficiency, not silicate. Together, the scenario of negative N*, positive Si*, and elevated Si:P portray a stoichiometric regime of acute NO limitation residual PO ³ and latent Si(OH_4_)^-1^, conditions that favor dominance of picocyanobacteria (Selph et al., 2015). Their small size and efficient uptake permit their persistence under intense light and scarce nutrients, (Flombaum and Martiny, 2021; Halsey et al., 2010, 2014; Six et al., 2021). Although ALFA does not distinguish between picocyanobacterial groups, the study of (Selph et al., this issue-a) which uses independent measurements from a flow cytometer, HPLC derived phytoplankton pigments including divinyl chlorophyll a (DV-Chl *a*), an exclusive marker pigment for *Prochlorococcus* and epifluorescence microscopy established that *Prochlorococcus* dominated the phytoplankton community. *Prochlorococcus* had the highest biomass of any phytoplankton group in all cycles while *Synechococcus* numbers remained unchanged and their biomass was a minor part of the total autotrophic biomass. Another interesting finding based on DNA and acid vacuole staining was that a considerable eukaryotic population was mixotrophic especially in Cycles 3 and 4, regions characterized by stronger stratification and warmer surface temperatures (Selph et al., this issue-b). Mixotrophy (Mitra et al., 2023), the ability to utilize both phototrophy and phagotrophy can become highly prevalent especially in oligotrophic, nutrient limited waters as this dual strategy augments nutrition (Schenone et al., 2024).

High resolution measurements of surface Chl *a* mirrored the spatial variability of the hydro-chemical environment detailed above. It was uniformly low, with a deep chlorophyll maximum (DCM) at about σθ ≈ 22.5–23 (≈26–27 °C), which shallowed and intensified whenever isopycnals domed due to eddy activity. Such intermittent events favored brief, localized increases in the cell counts and biomass of micro and nanoplankton (Fig 11), but their imprints were narrow, short-lived and transient (Fernández et al., this issue). In the lower euphotic zone, the aforementioned study of Selph et al. (this issue-a) characterized a diverse phytoplankton community of prymnesiophytes, dinoflagellates, prasinophytes, pelagophytes, and cryptophytes in addition to the ubiquitous *Prochlorococcus*.

### 4.2. Photophysiological insights

Spatial patterns of photo-physiological variables provided critical insight into the condition and functioning of phytoplankton in the oligotrophic seascape of the Argo Basin. Persistently low, surface maximum quantum yields of PSII (F_v_/F_m_ generally <0.5) were consistent with the chronic light stress and acute nitrate depletion that define this and other regions worldwide (Behrenfeld and Milligan, 2013; Gorbunov and Falkowski, 2020; Kranz et al., this issue; Oxborough et al., 2012; Schallenberg et al., 2020; Schuback and Tortell, 2019).

Turnover time τ_1_or the rate of primary electron transfer on the acceptor side of PSII can also provide additional functional diagnostics of nutrient stress. Short τ values can indicate fast electron transfer, typical of healthy, nutrient-replete phytoplankton with well-functioning PSII reaction centers. Longer τ values indicate a slower re-oxidation of Q, which can signal stress or limitation especially nitrogen or iron both of which restricts synthesis of electron transport proteins and PSII repair machinery. The connectivity factor (*p,* dimensionless) which is rarely exploited has been shown by Xu et al (2017) to be good indicator of photoinhibition. The factor _ρ_ measures excitation transfer from closed PSII to remaining open PSII upon illumination, which could theoretically generate a progressive increase in σPSII for the remaining open PSII. In laboratory experiments with 4 taxa including a culture of *Prochlorococcus*, with increasing actinic illumination PSII closed progressively and _ρ_ decreased. This was especially so with the *Prochlorococcus* which showed distinct rates of response of ρ to PSII closure from light exposure likely reflecting differences in the spacing or orientation among their PSII centers.

When a light limited *Prochlorococcus* culture was exposed to increasing light, *p* increased rapidly to about 0.8 at 800 *μ*mol photons m^-2^s^-1^ while a saturated light culture of the same rapidly decreased its *p* to 0.1. Our *p* values are akin to the latter indicating that phytoplankton in the ARGO Basin were in addition to nutrient limitation could also be photoinhibited.

σPSII, the functional absorption cross section of photosystem II, offers a dual perspective, one that relates to both the nutrient physiology and light acclimation as well as community composition. Smaller phytoplankton such as *Prochlorococcus* typically possess relatively small σPSII due to their reduced light-harvesting antennae whereas small eukaryotes such as cryptophytes generally exhibit larger cross-sections. The observed spatial distribution of σPSII, therefore, suggests that community composition varies subtly across the basin. Areas with higher σPSII values likely reflect a greater relative contribution picoeukaryotes, whereas areas of consistently low σPSII between 115° and 120°E and south of 15°S are most compatible with *Prochlorococcus* dominance. These inferences align with the cell count data reported by (Selph et al., this issue) and (Yingling et al., this issue), which show overwhelming *Prochlorococcus* numerical dominance punctuated by brief, localized increases in eukaryotic contribution as also seen in our phytoplankton community composition data collected during Cycles 1 to 4 (Figs. 12a-d) However, community composition changes are ‘overshadowed’ by the change of nutrient limitation and light acclimation status, both of which changed from east to west.

The photophysiological responses of picocyanobacteria reflected this stoichiometric setting. Cycle 1 followed a large tropical storm which deepened the mixed layer (Fig. S4a,e,f, m) and supplied some nitrate into surface waters. Cloud cover during the first 3 days prevented photoinhibition also. Diel patterns showed early morning highs, elastic midday depressions in PSII efficiency (F_v_/F_m_) that fully recovered overnight, consistent with a community supported by weak nutrient exchange due the interaction of the SICC with ITF, and capable of maintaining protein-rich PSII repair. The tight coupling between PAR and photophysiological properties is particularly evident on 6 February, a day marked by heavy cloud cover. During this period, lower and rapidly fluctuating PAR coincided with a shallower decline in σPSII and a rapid recovery of Fv/Fm values.

Cycle 2 exhibited several hydrographic features that distinguished it from the other cycles (Fig. S4a-p). Compared with Cycle 1, the mixed layer was much shallower during Cycle 2 and the upper ocean was much warmer than all the other Cycles. The surface waters were relatively fresh, suggesting the influence of lateral advection or freshwater input. The density field indicated a re-intensification of stratification, although gradients remained weaker than those observed in Cycles 3 and 4. Biologically it is clear that picocyanobacterial communities during Cycle 2 were nitrate stressed as their photophysiological responses were more variable and vulnerable to perturbations.

In Cycle 3, the physiology of the picocyanobacterial community also bore the imprint of chronic nitrate depletion: depressed predawn F_v_/F_m_ values, incomplete nightly recovery, reduced σPSII, and prolonged τ1 and τ2 electron turnover times. These patterns reflect the inability of cells to sustain nitrogen-intensive PSII repair cycles, leading to persistent downregulation. At this stage, the stoichiometric imbalance was stark, while both phosphate and silicate were present in relative excess, nitrogen was absent, restricting the community to high-light–adapted *Prochlorococcus* and *Synechococcus*. Cycle 4 which was closest to the shelf showed a different pattern with low muted diel patterns and an increase in PSII connectivity (*p*). We are unable to make any conclusions on differences of these turnover times across the 4 cycles and the increase in cycle 4 is specially confounding as other variable fluorescence parameters and (Kranz et al., this issue) indicate signs of photoinhibition for cells of this cycle. Net primary measurements by Kranz et al. this issue also revealed that average photosynthetic rates were lowest during Cycle 4.

Superimposed on this nitrogen-driven framework is the likely role of iron as a secondary limiting factor (Li et al., this issue) as indicated by the longer turn over times, *τ*_1_ and *τ*_2_ indicating a phytoplankton population that was acutely nitrogen as well as iron limited. Iron is required for electron transport proteins and PSII repair, and its scarcity can prolong electron turnover times (τ1 and τ2) and suppress F_v_/F_m_ even when macronutrients are partially relieved (Behrenfeld and Milligan, 2013; Schallenberg et al., 2020; Schuback et al., 2016; Schuback et al., 2015). The Argo Basin receives minimal atmospheric dust deposition compared to the Arabian Sea or northwest Indian Ocean (Guieu et al., 2019), limiting aeolian Fe inputs. Moreover, ITF waters entering the basin from the Pacific are Fe-poor (Pavia et al., 2020), and while continental shelf sediments can supply iron locally, the deep, stratified setting of the Argo Basin restricts vertical resupply. In this context, the combination of strongly negative N* and subtle Fe stress likely explains the persistent downregulation of PSII function observed across Cycles 3 and 4: nitrate scarcity limited protein-rich repair machinery, while iron scarcity constrained the cofactors required for efficient electron transport.

In synthesis, the transect-wide diel photophysiology of picocyanobacteria in the Argo Basin downstream of the ITF reflects a dynamic interplay between chronic nitrate and possible iron limitation, intense light forcing that can lead to photoinhibition, and episodic mesoscale variability. Negative N*, positive Si*, and high Si:P ratios indicate a nitrate-depleted but phosphate- and silicate-replete system with N_2_ fixation mitigating some nitrate limitation and supporting about 16% of the primary production. *Prochlorococcus* dominate through photophysiological adaptations by modulating light harvesting and nocturnal PSII repair to maintain productivity across a range of light conditions, despite nutrient limitation (Kranz et al., this issue). *Prochlorococcus* form the core of the community, sustaining the microbial food web and supporting higher trophic levels, while larger eukaryotes contribute episodically to biomass during nutrient relief events. This oligotrophic seascape is structured by the ITF–SICC frontal system, with a nutrient-poor eastern cap and a nutrient-accessible lower-euphotic frontal zone. Mesoscale eddies intermittently tilt density surfaces, granting transient nutrient pulses that briefly support larger phytoplankton and alter the depth and character of the DCM. These physical-biogeochemical couplings underscore how subtle shifts in frontal dynamics can drive disproportionate impacts on regional productivity and export.

This and other studies (Kranz et al., this issue; Landry et al., this issue-a; Landry et al., this issue-b; Landry et al., this issue-c; Selph et al., this issue-a; Selph et al., this issue-b; Stukel et al., this issue; Swalethorp et al., this issue; Yingling et al., this issue) that form BLOOFINZ project can provide highly significant insights into what makes the ARGO basin the optimal spawning habitat of SBT. Linear inverse ecosystem modeling (Stukel et al., this issue; Stukel et al., 2024) with assimilation of field data especially grazing experiments (Swalethorp et al., this issue) suggests a short and efficient food chain. Higher rates of herbivory were a combination of the high rates of feeding appendicularians on *Prochlorococcus* and crustaceans feeding on nanophytoplankton. In turn, grazing rates revealed appendicularians as the most significant and preferred prey of SBT larvae. These findings highlight the importance of transfer of energy via picocyanobacteria, and how a seemingly unproductive microbial base can underpin a globally significant pelagic ecosystems, the Argo basin of the tropical Indian Ocean. A recent paper (Ribalet et al., 2025) using a global ocean ecosystem model, shows that under future warming scenarios, there could be a possible 17–51% reduction in *Prochlorococcus* production in tropical oceans highlighting the potential vulnerability of this SBT spawning ground.

## 4. Conclusions

The slew of spatially high resolution photophysiological measurements undertaken in this study allowed us to understand how various phytoplankton populations respond to nutrient supplies in an oligotrophic seascape. Acute nutrient impoverishment (primarily nitrogen) is alleviated by N_2_ fixation from the large cyanobacterial populations, as well regenerated nitrogen and new nitrogen brought up by episodic mixing events. *Prochlorococcus* dominate through photophysiological adaptations by modulating light harvesting and nocturnal PSII repair to maintain productivity across a range of light conditions, despite nutrient limitation.

Our observations from the Argo Basin may offer a window into future ocean ecosystems, where climate-driven warming, stratification, and oligotrophication promote dominance by picocyanobacteria. Though small, these microbes efficiently channel energy through the food web, sustaining higher trophic levels, including apex predators such as bluefin tuna. Satellite records already show that oligotrophic regions are expanding and stratifying (Capotondi et al., 2012), favoring small-celled phytoplankton as primary producers (Fernández-González et al., 2022). Climate models predict that such conditions will intensify, and while picophytoplankton are projected to increase in abundance under warming scenarios (Fu et al., 2016; Peter and Sommer, 2013), although the recent study of Ribalet et al. (2025) based on decade-long field measurements shows that 17-51% reduction in *Prochlorococcus* in tropical oceans, which portends severe repercussions for this vital tuna spawning ground, the food chain of which is primarily supported by *Prochlorococcus*.

## Acknowledgements

The authors thank the captain and crew of the R/V *Roger Revelle* and the science team for their support and contributions during the field study. This study was supported by U.S. National Science Foundation Rapid Grant OCE-2332036 and NASA Grant 80NSSC20K0014 to JIG and HRG, and NSF grants OCE-2019983, 1851558 (to MRL) and 1851347 and 2404504 (to SAK).

Water and plankton samples were collected under Australian Government permit AU-COM2021-520 and Australian Marine Parks permit PA2021-00062-2 issued by the Director of National Parks, Australia. The views expressed in this publication do not necessarily represent those of the Director of National Parks or the Australian Government.

## Author Statement

JIG and HRG planned the field study and JIG conducted all experiments and prepared figures. HRG, and XW analyzed the data and prepared the figures, JW processed the satellite data. JIG deployed the ALFA, FIRe and FlowCAM instruments and conducted the field observations.

SAK provided the nutrient data. MRL planned the cruise and coordinated all sampling activities on board. RS helped process the FlowCAM data and calculation of carbon biomass. JIG and HRG wrote the manuscript, and all authors provided feedback on concepts in the manuscript, and provided edits of the manuscript.

## Supplementary figures

**Fig. S1.**
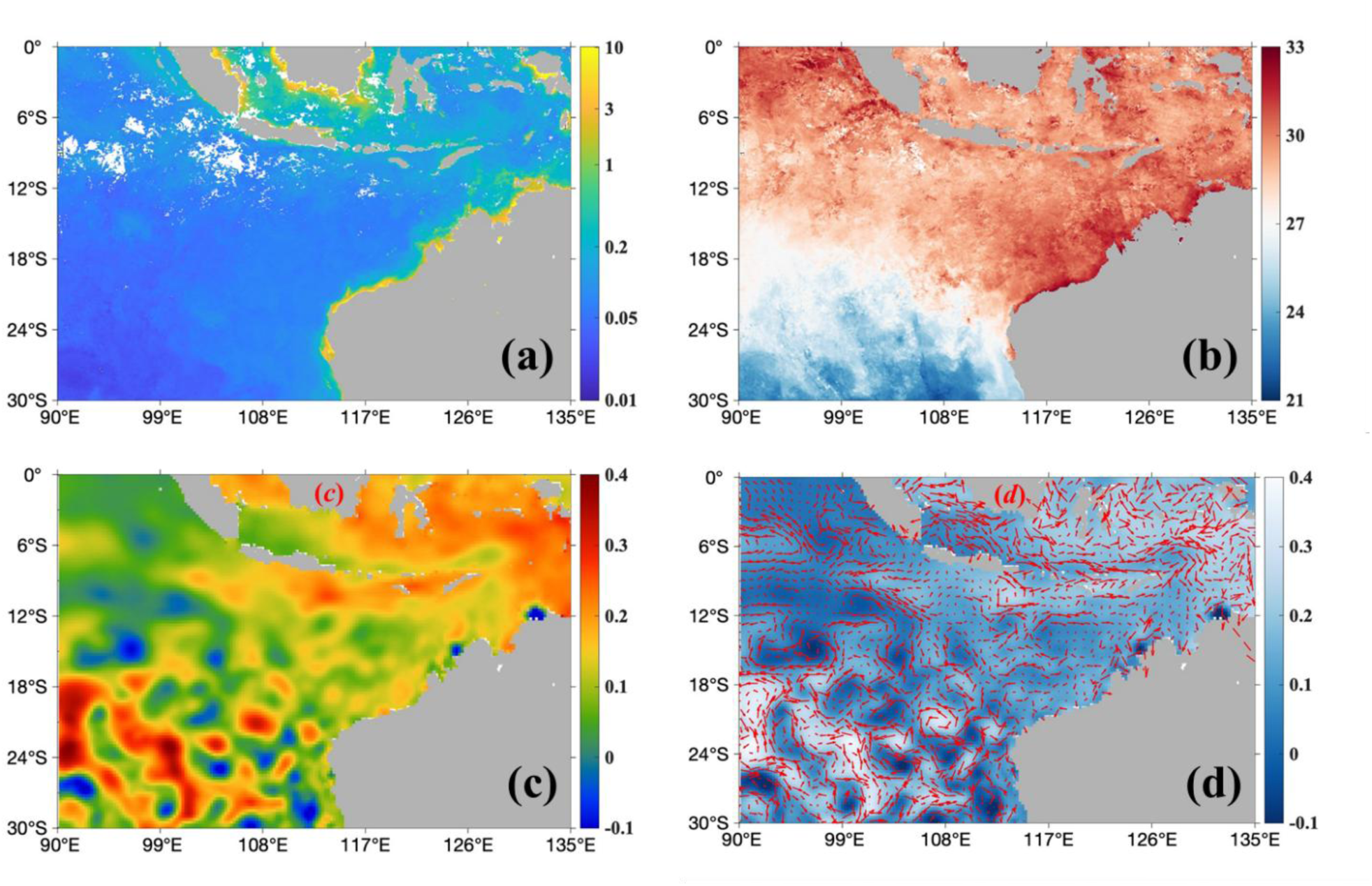
Monthly a) SNPP-VIIRS Sea surface Chlorophyll *a* (mg m^-3^) b) SNPP-VIIRS Sea surface temperature (°C) c) Aviso Sea Surface Height Anomaly (MSLA, m) and d) Aviso Sea surface current index FSLE (Finite-Size Lyapunov Exponents, day^-1^) from Copernicus Marine and Environment Monitoring Service.

**Fig. S2.**
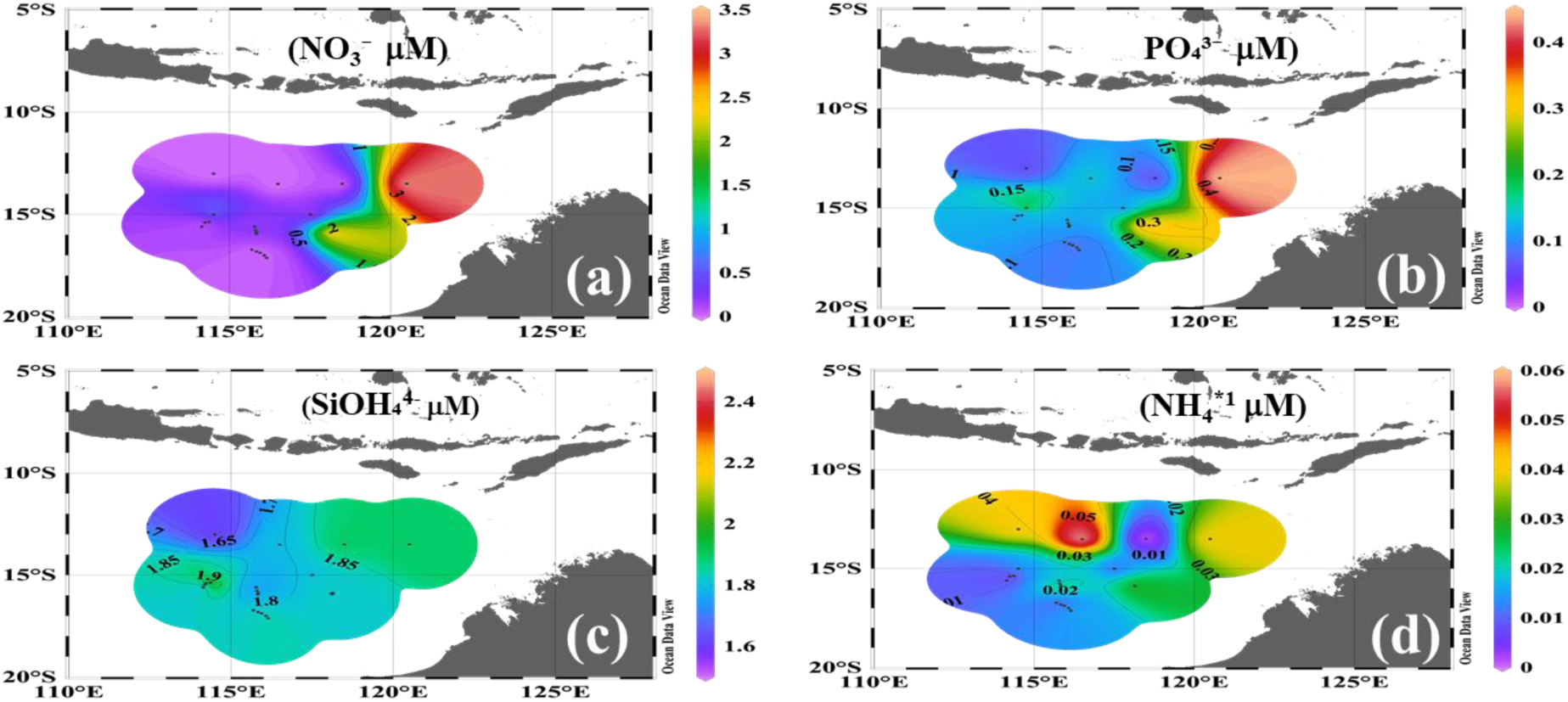
Spatial distribution of Nitrate (NO3^-^,μM), Phosphate (PO_4_^4-^ μM), Silicate (SiO_4_^4-^, μM) and Ammonium (NH^+^ μM) at 60 m in the Argo Basin. Other details as in Fig. 3

**Figure S3.**
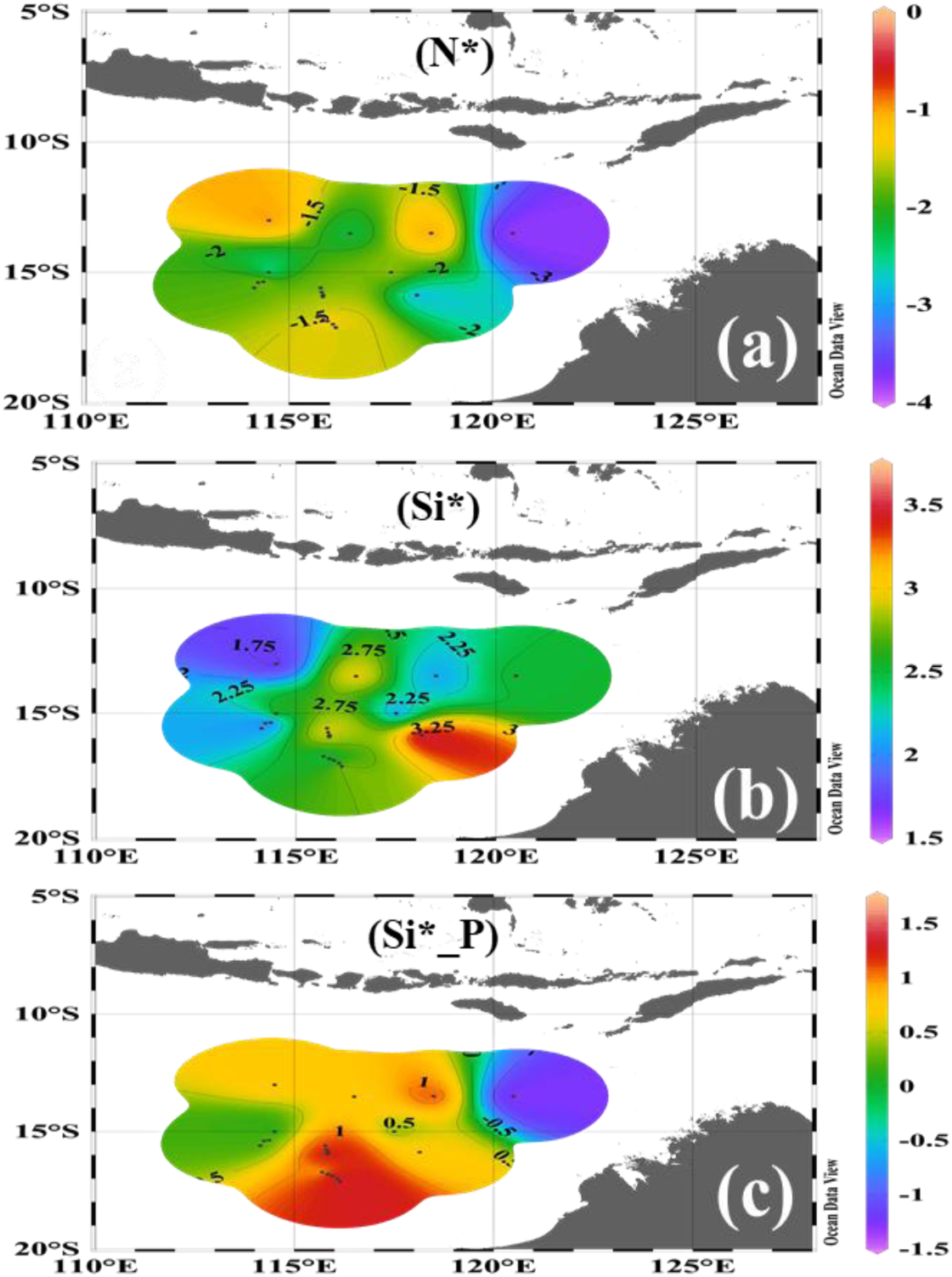
Spatial distribution of N*, Si* and Si:P at 20m in the Argo Basin. Other details as in Fig. 3

**Figure S4.**
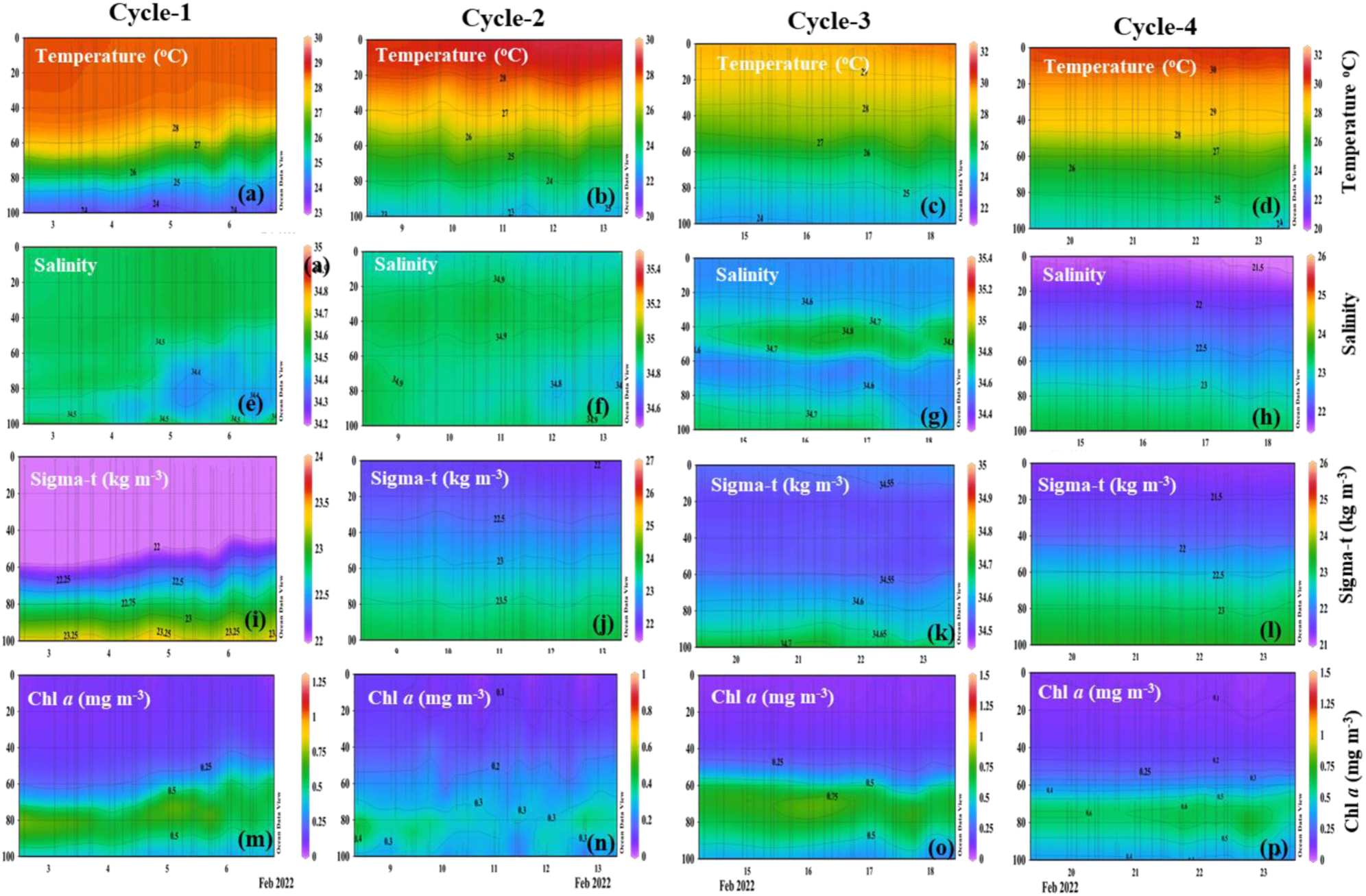
Hydrographic conditions during Cycles 1 to 4. Data were obtained from CTD deployments during the Cycle stations.

